# Mendelian randomization identifies folliculin expression as a mediator of diabetic retinopathy

**DOI:** 10.1101/2020.06.09.143164

**Authors:** Andrew D. Skol, Segun C. Jung, Ana Marija Sokovic, Siquan Chen, Sarah Fazal, Olukayode Sosina, Poulami Borkar, Amy Lin, Maria Sverdlov, Dingcai Cao, Anand Swaroop, Ionut Bebu, DCCT/ EDIC Study group, Barbara E. Stranger, Michael A. Grassi

**Affiliations:** Department of Pathology and Laboratory Medicine, Ann and Robert H. Lurie Children’s Hospital of Chicago, Chicago, IL; Research and Development, NeoGenomics Laboratories, Aliso Viejo, CA; University of Illinois at Chicago, Chicago IL; Cellular Screening Center, Office of Shared Research Facilities, The University of Chicago, Chicago, IL; Department of Biostatistics, Johns Hopkins University, Baltimore, MD; National Eye Institute, National Institutes of Health (NIH), Bethesda, MD; The George Washington University, Biostatistics Center, Rockville, MD; Department of Pharmacology, Center for Genetic Medicine, Northwestern University Feinberg School of Medicine. Chicago IL

## Abstract

The goal of the study was to identify genes whose aberrant expression can contribute to diabetic retinopathy. We determined differential gene expression in response to high glucose in lymphoblastoid cell lines derived from matched individuals with type 1 diabetes (T1D) with and without retinopathy. Those genes exhibiting the largest difference in glucose response between individuals with diabetes with and without retinopathy were assessed for association to diabetic retinopathy utilizing genotype data from a genome-wide association study meta-analysis. All genetic variants associated with gene expression (expression Quantitative Trait Loci, eQTLs) of the glucose response genes were tested for association with diabetic retinopathy. We detected an enrichment of the eQTLs from the glucose response genes among small association p-values and identified folliculin (*FLCN*) as a susceptibility gene for diabetic retinopathy. We show that expression of *FLCN* in response to glucose was greater in individuals with diabetic retinopathy compared to individuals with diabetes without retinopathy. Three large, independent cohorts of individuals with diabetes revealed an association of *FLCN* eQTLs to diabetic retinopathy. Mendelian randomization further confirmed a direct positive effect of increased *FLCN* expression on retinopathy in individuals with diabetes. Together, our studies integrating genetic association and gene expression implicate *FLCN* as a disease gene for diabetic retinopathy.

## Introduction

Almost all individuals with diabetes will develop some form of diabetic retinopathy over time [1]. In the United States diabetic retinopathy is the most frequent cause of blindness among working age individuals [2]. Interindividual variation contributes significantly to susceptibility of the severe manifestations of diabetic retinopathy, which results in vision impairment. Epidemiological studies suggest that phenotypic variation is influenced by two primary risk factors: the duration of diabetes, and an individual’s level of glycemia (HbA1c) [3]. However, these two factors do not completely explain an individual’s risk for developing diabetic retinopathy. For instance, a common anecdotal clinical experience is the comparison of patients with similar durations of diabetes and similar levels of glycemic control who have tremendously disparate clinical outcomes for diabetic retinopathy. Moreover, some individuals with diabetes develop very minimal retinopathy [4], whereas others clearly seem to have a predisposition for severe retinopathy [5].

Together, these observations in conjunction with the high concordance of diabetic retinopathy between family members support an underlying genetic mechanism. Familial aggregation and twin studies estimate that genetic factors account for 25 to 50 percent of an individual’s risk of developing severe diabetic retinopathy [6] [7]. Unfortunately, little is known about the genetic architecture that contributes to susceptibility for diabetic retinopathy. Genetic studies suggest that it is a highly polygenic trait influenced by multiple genetic variants of small effect. Our group and others have performed genomewide association studies to better delineate the molecular factors that predispose to diabetic retinopathy [8] [9] [10]. However, these studies have had limited success, likely due to insufficient study sample sizes and the phenotypic heterogeneity of diabetic retinopathy.

Notably, like other complex disease traits including age-related macular degeneration [11, 12], a majority of genetic variants nominally associated with diabetic retinopathy are located in intronic or inter-genic regions [13]. Most of these variants appear to play critically important functional roles in regulating gene expression. In fact, several of the top associated SNPs identified in our meta-GWAS of diabetic retinopathy [8] are present in DNase hypersensitivity sites and affect gene expression levels by altering the allelic chromatin state or the binding sites of transcription factors [14].

The observation that disease-associated genetic loci often influence gene expression levels [15] led us to postulate that integrating gene expression with genetic association would be a powerful approach to identify susceptibility genes for diabetic retinopathy. We hypothesized that cell lines derived from individuals with diabetes with and without retinopathy could be used to uncover genetic variation that explain individual differences in the response to diabetes. Culturing two sets of cell lines under controlled, identical conditions from individuals with diabetes who did and those who did not develop retinopathy, could unmask molecular differences in how these groups respond to glucose [16] [17]. We presumed that a portion of those differences would have a genetic basis.

In this article, we identify genes whose expression responds differently to glucose in cells derived from T1D individuals with and without diabetic retinopathy. We show that one of these genes, folliculin (*FLCN*), is causally implicated in diabetic retinopathy based on results from genetic association testing and Mendelian randomization.

## Methods

### Overview

In this study we profiled the transcriptomes of cell lines derived from 22 individuals [7 individuals with no diabetes (nDM), 8 with T1D with proliferative diabetic retinopathy (PDR) and 7 with T1D with no retinopathy (nDR)] utilizing gene expression microarrays to characterize the transcriptional response to glucose. Specifically, we cultured lymphoblastoid cell lines (LCLs) derived from each individual in standard glucose (SG) and high glucose (HG) medium and measured gene expression for each gene in each sample, as well as the difference (Δ = response to glucose [RG]) in each gene’s expression for each individual (Figure 1a). In this manuscript, ‘**Differential expression**’ refers to gene expression comparisons between groups (nDM, PDR, nDR) in the same condition (SG or HG) and between conditions (SG and HG) in the same group (nDM, PDR or nDR).

**Figure 1.**
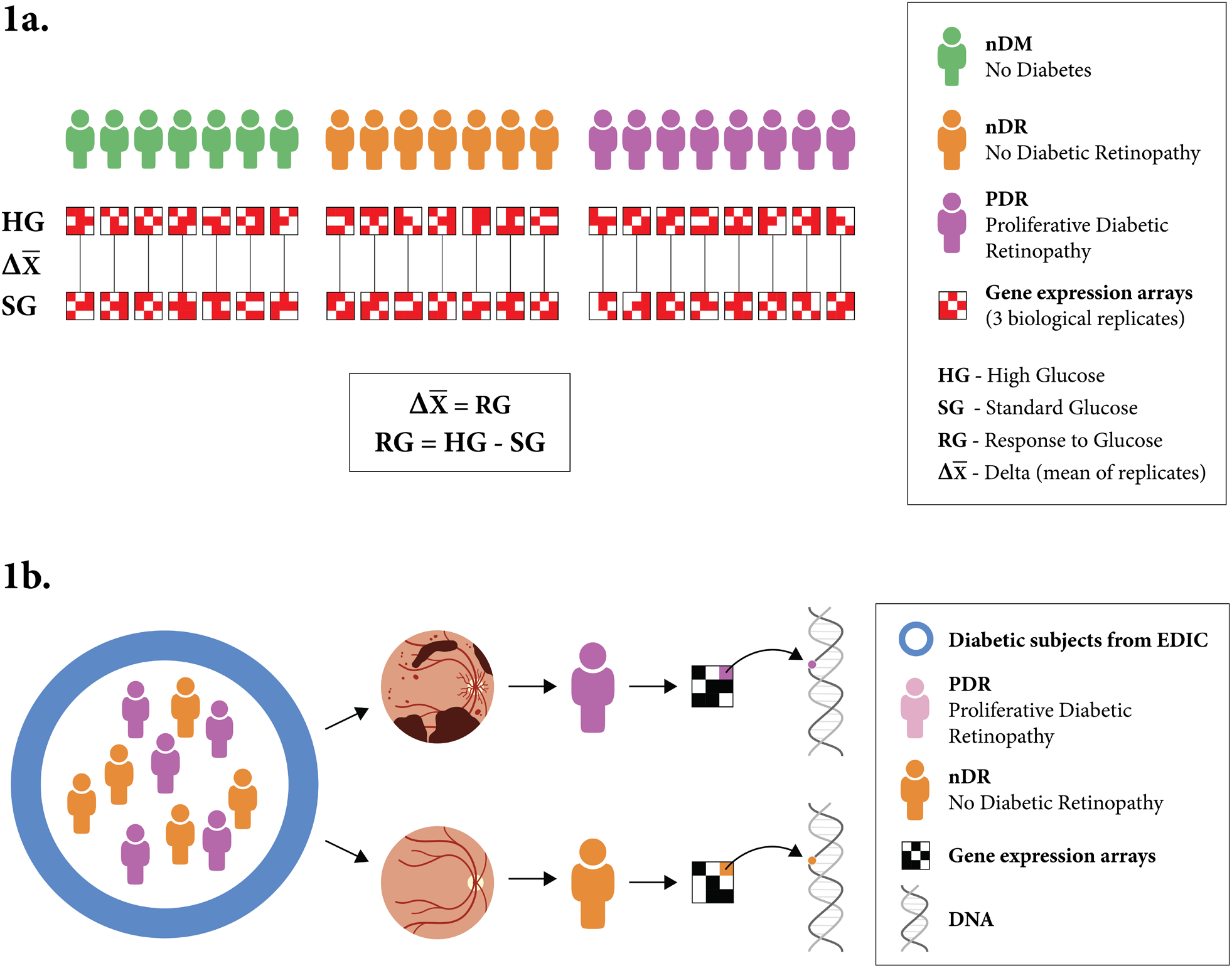
Experimental design. a) Schematic representation of the experimental design for transcriptomic profiling. Lymphoblastoid cell lines (LCLs) from 22 individuals were cultured under both standard glucose and high glucose conditions. Gene expression was quantified using microarrays for three technical replicates of each LCL in each condition. The response to glucose was determined for all genes on a per-individual basis, by comparing expression in standard and high glucose conditions. The cell lines were derived from individuals with diabetes and no retinopathy (7), individuals with diabetes and proliferative diabetic retinopathy (8), and individuals without diabetes (7). b) We identified 15 individuals based on retinopathy status from the Epidemiology of Diabetes Interventions and Complications (EDIC) cohort. We compared the differential response in gene expression to glucose for individuals with and without proliferative retinopathy (RG_pdr-ndr_). Expression quantitative trait loci (eQTL) for those genes that showed the greatest differential response between individuals with and without retinopathy were tested for their genetic association to diabetic retinopathy.

We compared the differential response in gene expression to glucose for all individuals with and without proliferative retinopathy. ‘**Differential response**’ in gene expression refers to the difference in gene expression response to glucose between groups. Specifically, we identified genes with a significant differential response in expression between individuals with diabetes with and without proliferative diabetic retinopathy (RGpdr-ndr).

We followed up genes showing differential response using the results of both a prior genome-wide association study meta-analysis of diabetic retinopathy (in the GoKinD and EDIC cohorts) [8] and the results of a multi-tissue expression quantitative trait loci (eQTL) analysis from GTEx [18] to identify potential diabetic retinopathy susceptibility genes (Figure 1b).

### Subject Safety and Confidentiality Issues

All cell lines were de-identified prior to their arrival at the University of Illinois at Chicago; therefore, this proposal qualified as non-human subjects research according to the guidelines set forth by the institutional review board at the University of Illinois at Chicago. As the data were analyzed anonymously, no subject consent was required. DCCT subjects previously provided consent for their samples to be used for research.

Matching of subjects was performed at George Washington University Biostatistics Center and did not involve protected health information as the phenotypic data were deidentified. The George Washington University institutional review board has approved all analyses of EDIC data of this nature. All protocols used for this portion of the study are in accordance with federal regulations and the principles expressed in the Declaration of Helsinki. Specific approval of the study design and plan was obtained from the EDIC Research Review Committee.

### Cell Lines

Twenty-two lymphoblastoid cell lines were used in the study as described previously [16]. Briefly, we included 15 of the 1,441 lymphoblastoid cell lines generated from individuals with type 1 diabetes from the DCCT/ EDIC cohort [19] [20], consisting of 8 matched cases with PDR (PDR) and 7 without retinopathy (nDR) as the controls [16] (Supplemental Table S1). Whole blood samples were ascertained from DCCT study subjects between 1991 and 1993. White blood cells from the samples were transformed into lymphoblastoid cell lines in the early 2000s. The fifteen lymphoblastoid cell lines from individuals with diabetes consisted of matched cases and controls. Cases were defined by the development of proliferative diabetic retinopathy by EDIC Year 10 (2004), whereas controls had no retinopathy through EDIC Year 10 (2004). Retinopathy grade was determined by seven-field stereoscopic photos. Control subjects had an ETDRS (Early Treatment Diabetes Retinopathy Score) of 10 and case subjects had an ETDRS score of ≥ 61. Most pairs were matched by age, sex, treatment group (intensive vs. conventional), cohort (primary vs. secondary), and diabetes duration [21] [22], except one pair that was matched by age, sex and treatment group only. Diabetes duration was defined as the number of months since the diagnosis of diabetes at DCCT baseline which was the time at subject enrollment (1983–1989). The remaining seven lymphoblastoid cell lines were purchased from the Coriell Institute for Medical Research NIGMS Human Genetic Cell Repository (http://ccr.coriell.org/) (GM14581, GM14569, GM14381, GM07012, GM14520, GM11985, and GM07344). None of these individuals had a history of diabetes (nDM). The covariates available for these 7 individuals were age and sex; male and female individuals were included. All of these individuals without diabetes were unrelated and of European ancestry [16] [17] (Supplemental Table S2).

### Culture Conditions

All lymphoblastoid cell lines were maintained in conventional lymphocyte cell culture conditions of RPMI 1640 with 10% FBS in a 25-cm^2^ cell culture flask. The cells were incubated at 37^0^C in 5% CO_2_ and the media was changed twice each week. Prior to the experiments (below), lymphoblastoid cells following serum starvation were passaged for a minimum of one week in either standard (SG) RPMI 1640 (11mM glucose) or high glucose (HG) RPMI media (30mM glucose) [23].

### Gene Expression Profiling

Quality control from RNA extraction was performed using the Agilent bio-analyzer, processed using the Illumina^™^ TotalPrep^™^-96 RNA Amplification Kit (ThermoFisher 4393543), hybridized to Illumina HT12v4 microarrays (Catalog number: 4393543), and scanned on an Illumina HiScan scanner [24] [25]. For each of the 22 individuals, three biological replicates were profiled, with each sample assessed at both standard glucose conditions (11mM of glucose), as well as high glucose conditions (30mM of glucose). Biological replicates were split from the same mother flask; cells were grown in separate flasks and run on different microarray plates on different days. Each biological replicate was generated from a separate frozen aliquot of that cell line. The gene expression profiling was performed in a masked fashion for both the case/control (PDR, nDR, nDM) status of the individual as well as the glucose treatment (SG, HG) of the sample in order to reduce any bias.

### Relative EBV Copy Number

Standard TaqMan qPCR was performed using EBV and NRF1 probes and primers [26]. To calculate real-time PCR efficiencies a standard curve of ten points of 2-fold dilution of 156.7 ng of gDNA was used from the Raji cell line (ATCC_®_ CCL-86^™^). Probes were designed for the target, EBV, and a reference gene, *NRF1*. Final concentrations of the probes and primers were 657nM and 250nM respectively. EBV probe: 5’6FAM-CCACCTCCACGTGGATCACGA-MGBNFGQ3’; EBV forward primer: 5’ GAGCGATCTTGGCAATCTCT; EBV reverse primer: 5’ AGTAGCCAGGCACCTACTGG; NRF1 probe: 5’VIC-CACTGCATGTGCTTCTATGGTAGCCA-MGBNFQ3’; NRF1 forward primer: 5’ ATGGAGGAACACGGAGTGAC; NRF1 reverse primer: 5’ CATCAGCTGCTGTGGAGTTG. Cycle number of crossing point versus DNA concentration were plotted to calculate the slope. The real-time PCR efficiency (E) was calculated according to the equation: E = 10 ^(−1/slope)^. Triplicates were done for each data point. Genomic DNA (78.3 ng) from each lymphoblastoid cell line was used in a standard TaqMan qPCR reaction with *EBV* as target gene and *NRF1* as reference gene. The sequences and concentrations of the probes and primers were as shown above.

### Growth Rate Measurement

Lymphoblastoid cell lines were thawed and cultured in RPMI and 10% FBS until they reached over 85% cell viability. Cells were seeded in a T25 flask. Two replicates were performed per cell line. Cells were counted every day or every other day for five to ten days and recorded.

### Quality Control for Gene Expression

The gene expression data comprised a total of 144 samples from 22 individuals (3 replicates per individual and treatment, except for 3 individuals with 5 replicates). Gene expression was assessed in two conditions, standard glucose and high glucose, and generated in four batch runs that were carefully designed to minimize potential batch effects. BeadChip data were extracted using GenomeStudio (version GSGX 1.9.0) and the raw expression and control probe data from the four different batches were preprocessed using a lumiExpresso function in the lumi R package version 2.38.0 [8, 9] in three steps: (i) background correction (lumiB function with the bgAdjust method); (ii) variance stabilizing transformation (lumiT function with the log2 option); (iii) normalization (lumiN function with the robust spline normalization (rsn) algorithm that is a mixture of quantile and loess normalization). To remove unexpressed probes, we applied a detection filter to retain probes with strong true signal by applying Illumina BeadArrays detection p-values < 0.01 followed by removing probes that did not have annotated genes, resulting in a total of 15,591 probes.

### Gene Expression Analysis

The study design is portrayed in Figure 1a. For a given individual S_i_ (*i*= 1,…,22) and gene G_k_ (*k*= 1,…,15591), we calculated Δ_i,k_= HG_i,k_ - SG_i,k_, where Δ is the individual’s response to glucose (RG), HG is gene expression in high glucose culture, and SG is gene expression in standard glucose culture. All replicate data were fit using a mixed model that accounted for the correlation between repeated measures within individuals.

The design matrix was constructed and analysis performed using the R version 3.5.1 package *limma* [27]. We built a design matrix using the model.matrix function, and accounted for correlation between biological triplicates using limma’s *duplicatecorrelation* function. A mixed linear model was then fit that incorporates this correlation and Δ_i,k_ using the *lmFit* function. Principal component analysis (PCA) of gene expression was run with the *prcomp* function in R [28]. For each gene, we calculated moderated t- and f-statistics and log-odds of expression by empirical Bayes moderation of the standard errors towards a common value. Differential expression is described using fold change (FC) while differential response reflects fold change (FC) differences between groups. The power to detect a 2 FC in gene expression between the two retinopathy groups (retinopathy vs. no retinopathy) (RGp_dr-ndr_)) given our sample size and using a type I error rate of 0.05 is 95% (as supported by our prior work [16])

### Gene set enrichment analysis (GSEA)

GSEA was performed using pre-ranked gene lists [29]. We ranked all analyzed genes based on sign (fold change) x (−log_10_(p-value)) [9]. Duplicated genes were removed. The gene ranking resulted in the inclusion of 11,579 genes. Enrichment statistics were calculated using rank weighting and the significance of enrichment was determined using permutations performed by gene set. The gene sets included c2.all.v6.0 and c5.all.v6.0. The minimum gene set size was 15 and the maximum gene set size was 500. GSEA was used to identify significant gene sets for the response to glucose in all study subjects (RG_all_ : nDM + PDR + nDR).

### Expression quantitative trait loci (eQTL)

To determine if the genes showing a differential response in gene expression (RG_pdr-ndr_) is driven by germline genetic variation, we tested if the eQTLs for these genes are enriched for small diabetic retinopathy GWAS p-values [8]. We use the term, ‘differential response gene’, for those genes identified by the RG_pdr-ndr_ analysis. All statistically significant eSNPs (false discovery rate (FDR) threshold of ≤0.05) (single nucleotide polymorphisms, SNPs, corresponding to *cis*-eQTLs from the GTEx and EyeGEx datasets) were collated for the glucose response genes in any of the 48 GTEx (version 7) tissues and the retina [18] [30]. We use the term eGene for any gene with at least one significant eSNP in any tissue.

### Genome-wide association study (GWAS)

Meta-analysis p-values were ascertained from our prior genome-wide association study for diabetic retinopathy [8]. The study assessed the genetic risk of sight threatening complications of diabetic retinopathy as defined by the presence of diabetic macular edema or proliferative diabetic retinopathy (cases) compared to those without (controls) in two large type 1 diabetes cohorts of 2829 total individuals (973 cases, 1856 controls) taken from the Genetics of Kidney in Diabetes (GoKinD) and the Epidemiology of Diabetes Interventions and Complications study (EDIC) cohorts.

We sought to determine whether there is enrichment of small p-values for diabetic retinopathy meta-GWAS among the significant eQTLs for the glucose response genes that show a significant differential glucose response between individuals with and without retinopathy (RG_pdr-ndr_). We used Benjamini-Hochberg adjusted p-values (FDR) to account for multiple testing given the high level of linkage disequilibrium between many eSNPs within an eQTL. SNPs from the three studies (expression, eQTL, GWAS) were matched by mapping all SNPs to dbSNP v.147 [31]. We determined the corresponding FDR for each glucose response gene’s eSNPs in the diabetic retinopathy meta-GWAS. To assess enrichment, we first determined the observed proportion of meta-GWAS FDR values < 0.05 among the statistically significant eQTLs of the glucose response genes (RG_pdr-ndr_). Next, we took 10,000 random samples of 103 GTEx eGenes (genes with an eQTL in any GTEx tissue) and identified corresponding eSNPs across all GTEx tissues. We calculated the GWAS FDR for these eSNPs and recorded the proportion of FDR values < 0.05.

Validation for the association of glucose response gene eSNPs with diabetic retinopathy was performed in the UK Biobank (UKBB) GWAS (Supplemental Table S3) [32]. Only individuals of northern European ancestry were analyzed. Quality control excluded individuals who were outliers based on relatedness; exhibited an excess of missing genotype calls; had more heterozygosity than expected; or had sex chromosome aneuploidy. A total of 337,147 individuals were available for analysis. Case subjects were defined as those who answered “yes” to questionnaire data eyesight field 6148 ‘Diabetes related eye disease’ (n=2,332). Our prior work validated the utility of self-report for the presence of severe diabetic retinopathy [31] [33]. Control subjects were defined as those who answered “yes” to data field 2443 ‘Diabetes diagnosed by doctor’ (n=14,680), excluding case subjects. SNPs were excluded according to the following: minor allele frequency < 0.004; missing rate > 0.015; HWE p-value < 1×10^−10^; INFO score < 0.8. We performed logistic regression as implemented in Plink2 [34] on this set of cases and controls. The logistic regression, including the following covariates: first 10 genotype-based principal components; chromosomal sex (as defined by XX, XY status); age; type of diabetes; HbA1c; and genotyping array type.

### Mendelian Randomization

To explore the causal effect of increased folliculin (*FLCN*) expression on diabetic retinopathy, we employed Mendelian randomization [35]. Effects were estimated with summary data-based Mendelian randomization analysis [36] (SMR). We estimated the effect of increasing levels of *FLCN* expression on diabetic retinopathy in the UKBB GWAS for diabetic retinopathy (described above) utilizing 272 SNPs that were significant cis-eSNPs (FDR ≤0.05) for *FLCN* in retina and also in at least 20 GTEx tissues. A total of 246 SNPs remained after removing those SNPs or their proxies (r^2^#x003E; 0.8) not genotyped in the UKBB. For each individual, the exposure was based on the genetically predicted gene expression of *FLCN* in retina and the outcome was the likelihood of having diabetic retinopathy. Heterogeneity in dependent instruments (HEIDI) [36] was used to investigate the possibility of confounding bias from horizontal pleiotropy with 14 independent (r^2^ < 0.2) *FLCN* eQTLs. As multiple independent (r^2^ < 0.2, n = 14) *FLCN* eQTLs exist, we also employed multi-SNP Mendelian randomization to assess for an aggregated effect [37] of the eQTLs on diabetic retinopathy mediated through *FLCN* expression.

### Folliculin (FLCN) Localization in Human Donor Eye Retina

A whole eye from a 69-year old Caucasian female post-mortem donor without diabetes was obtained from National Disease Research Interchange (NDRI). Findings were replicated in an additional five post-mortem donors without diabetes from the NDRI. The eye was cut in half in a horizontal plane, and each half was placed in an individual cassette. Samples were processed on ASP300 S automated tissue processor (Leica Biosystems) using a standard overnight processing protocol and embedded into paraffin blocks. Tissue was sectioned at 5 μm, and sections were de-paraffinized and stained on BOND RX autostainer (Leica Biosystems) following a preset protocol. In brief, sections were subjected to EDTA-based (BOND ER2 solution, pH9) antigen retrieval for 40 min at 100°C, washed and incubated with protein block (Background Sniper, Biocare Medical, BS966) for 30 min at room temperature. For immunofluorescence (IF), sequential staining with rabbit polyclonal anti-FLCN antibody (1:50, Abcam #ab93196) and mouse monoclonal anti-CD31 antibody (1:50, DAKO, M0823) was conducted using goat-anti-rabbit Alexa-488 and goat-anti-mouse Alexa-555 secondary antibodies (Molecular Probes) for detection. DAPI (Invitrogen, #D3571) was used to stain nuclei. The slides were mounted with ProLong Diamond Antifade mounting media (ThermoFisher, #P36961). Images were taken at 20x magnification on Vectra 3 multispectral imaging system (Akoya Biosciences). A spectral library acquired from mono stains for each fluorophore (Alexa-488, Alexa-594), DAPI, and human retina background fluorescence slide was used to spectrally unmix images in InForm software (Akoya Biosciences) for visualization of each color.

### Data and Code Availability

The microarray expression data are available at Gene Expression Omnibus (GEO) under accession code GSE146615. The cleaned analysis dataset of the diabetic retinopathy GWAS in the UKBB will be uploaded to the UKBB archive (https://oxfile.ox.ac.uk/oxfile/work/extBox?id=825146B4380F72048D). Please contact for further information.

## Results

### Individuals with retinopathy (PDR) show differences in diabetes duration and level of glycemia compared to individuals without retinopathy (nDR)

Matched DCCT/EDIC subjects from whom the gene expression profiling was obtained are detailed in Supplemental Table S1. All individuals had T1D, were Caucasian, and 60% were female. As anticipated, notable differences were observed between individuals with and without retinopathy (PDR vs. nDR) for duration of diabetes (53 +/43.4 months vs. 27 +/- 13.4 months) and mean HbA1c (9.71 +/- 2.37 vs. 7.62 +/- 1.07), respectively, given their significant impact on retinopathy.

### Interindividual variation is evident in the transcriptional response to glucose

We quantified gene expression levels from LCLs of all study individuals (nDM, PDR, nDR) in both standard glucose (SG) and high glucose conditions (HG) and determined the genome-wide transcriptional response to glucose for each individual (RG_all_). We observed that 22% of 11,548 examined genes were differentially expressed between the two conditions (true positive rate; π_1_ = 0.22) [38] (Supplemental Figure S1), with 299 of those at an FDR < 0.05 (Figure 2a), supporting a significant impact of glucose on the LCL transcriptome. We confirmed that interindividual transcriptome response to high glucose is greater than the intraindividual response (P = 2 x 10^−16^) (Supplemental Figure S2a-c). Interestingly, *TXNIP*, the most highly glucose-inducible gene in multiple cell types [39, 40], exhibited the largest (log_2_(FC) difference = 0.2) and most significant (P = 3.2 ×10^−12^, FDR = 5.1 ×10^−8^) transcriptional response to glucose. Pathway analysis using Gene Set Enrichment Analysis (GSEA) revealed dramatic up-regulation of genes involved in structural changes to DNA (DNA packaging, FDR < 0.0001; nuclear nucleosome, FDR = 0.001) and in genes such as transcription factors that modulate the cellular response to environmental stimuli (protein DNA complex, FDR < 0.0001) (Figure 2b). Conversely, genes that modulate the cellular response to infection were considerably down-regulated (type 1 Interferon, FDR < 0.0001; gamma Interferon, FDR < 0.0001; leukocyte chemotaxis genes, FDR < 0.0001). This finding suggests that chronic glucose exposure depresses cellular immune responsiveness and may explain in part the increased risk of infection found in patients with diabetes [41] [42].

**Figure 2.**
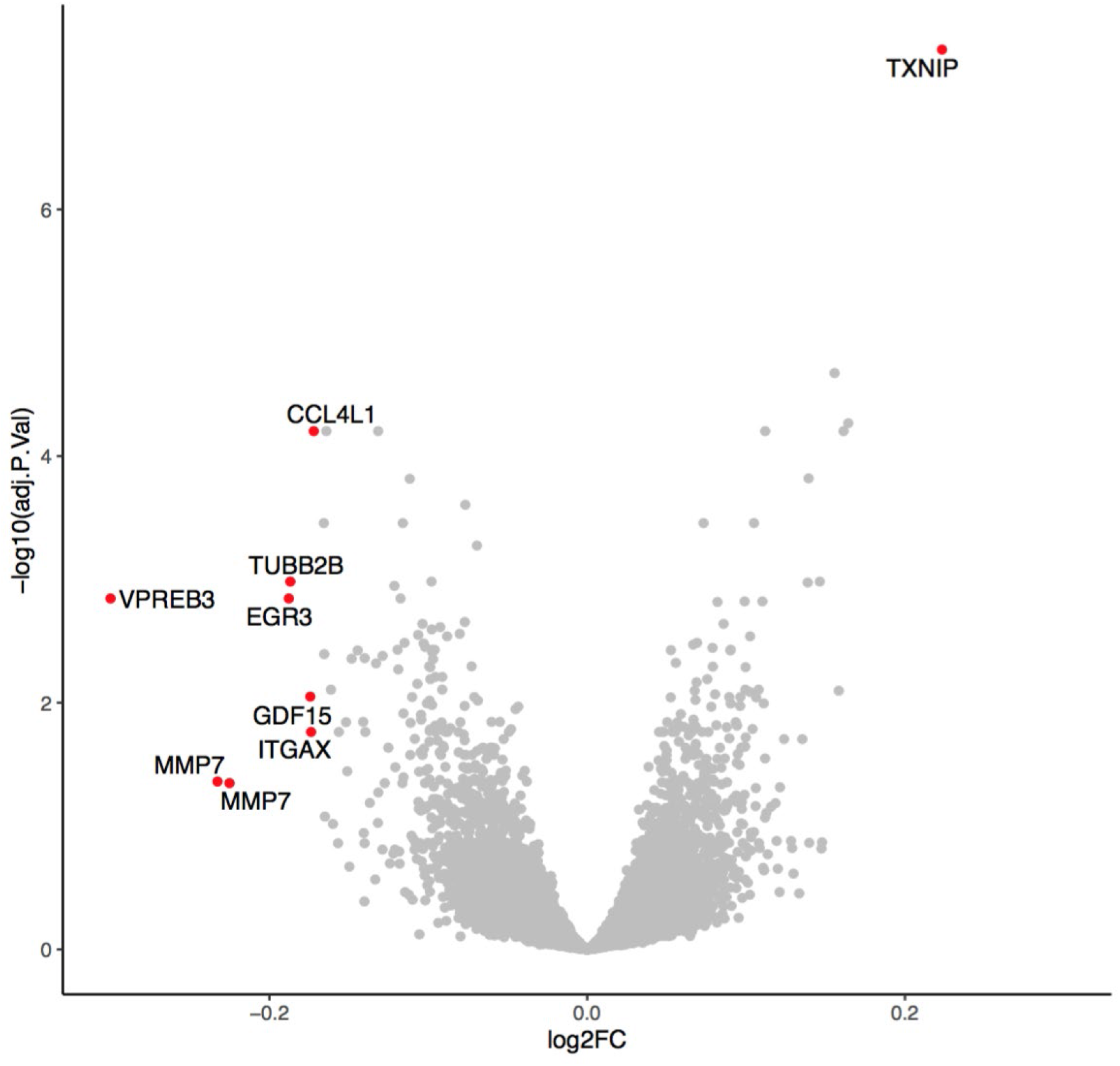
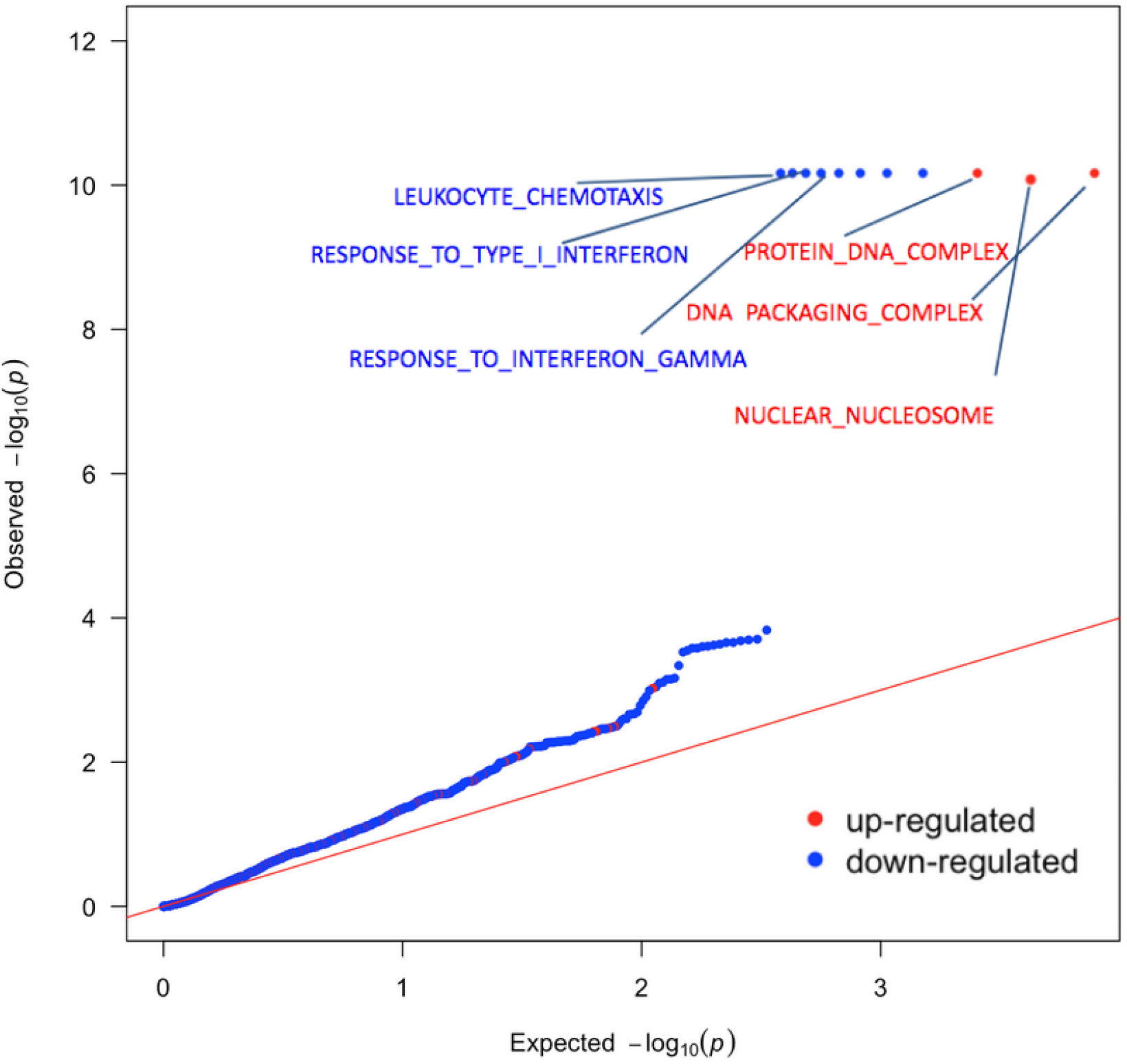
Response to glucose. a) Volcano plot summarizing transcriptional response to glucose for all 22 individuals (RG_All_ consisting of nDM, nDR and PDR individuals). Each point represents a single gene. Red indicates differentially expressed genes (FDR < 0.05 (log_10_ > 1.3 represented by the dotted line) and an absolute log_2_FC > 0.17. Adj p-value is false discovery rate (FDR). FC indicates expression fold change with positive values indicating higher expression in the high glucose condition relative to the standard condition. b) QQ plot summarizing GSEA of transcriptional response to glucose in all 22 individuals. Pathways are classified as up-regulated (red) or down-regulated (blue) in response to glucose. Only significant GO categories (FDR < 0.1%) are labeled. Red line indicates the null expectation.

### Individuals with diabetic retinopathy exhibit a differential transcriptional response to glucose

We observed differences in the transcriptional response to glucose between matched individuals with and without diabetic retinopathy (RG_pdr-n_d_r_). Principal component analysis (PCA) demonstrated that the observed interindividual variance is dominated by randomized DCCT treatment (intensive vs. conventional) group effects based on retinopathy status (P = 3 x 10^−6^) (Supplemental Figure S3) and is not confounded by LCL growth rate (P > 0.05) or EBV copy number (P > 0.05). Using a gene-wise analysis we identified 19 genes exhibiting a differential glucose response between individuals with and without retinopathy (P < 0.05, absolute log_2_ FC difference > 0.26) (Figure 3; Supplemental Table S4). Some of these genes and pathways have previously been shown to play a role in diabetic retinopathy. One of the top differential response genes was *IL1B* (P = 0.008, log_2_(FC) response difference = 0.289). Expression of *IL1B* has been previously reported to be induced by high glucose [43]. Additionally, the expression of *IL1B* is upregulated in the diabetic retina and has been implicated in the pathogenesis of diabetic retinopathy [44]. Likewise, the top GSEA pathway has also previously been implicated in the pathogenesis of diabetic retinopathy. We identified PDGF signaling as the most significant differential response pathway (FDR = 0.012) (Supplemental Figure S4). Elevated levels of PDGF are present in the vitreous of individuals with proliferative diabetic retinopathy compared to individuals without diabetes [45]. As PDGF is required for normal blood vessel maintenance, it is thought to contribute to the pericyte loss, microaneurysms, and acellular capillaries that are key features of the diabetic retina [46]. Interestingly, despite our model utilizing lymphoblastoid cells, it was able to reveal the upregulation of PDGF which is primarily a vascular factor that also plays a key role in neuronal tissue.

**Figure 3.**
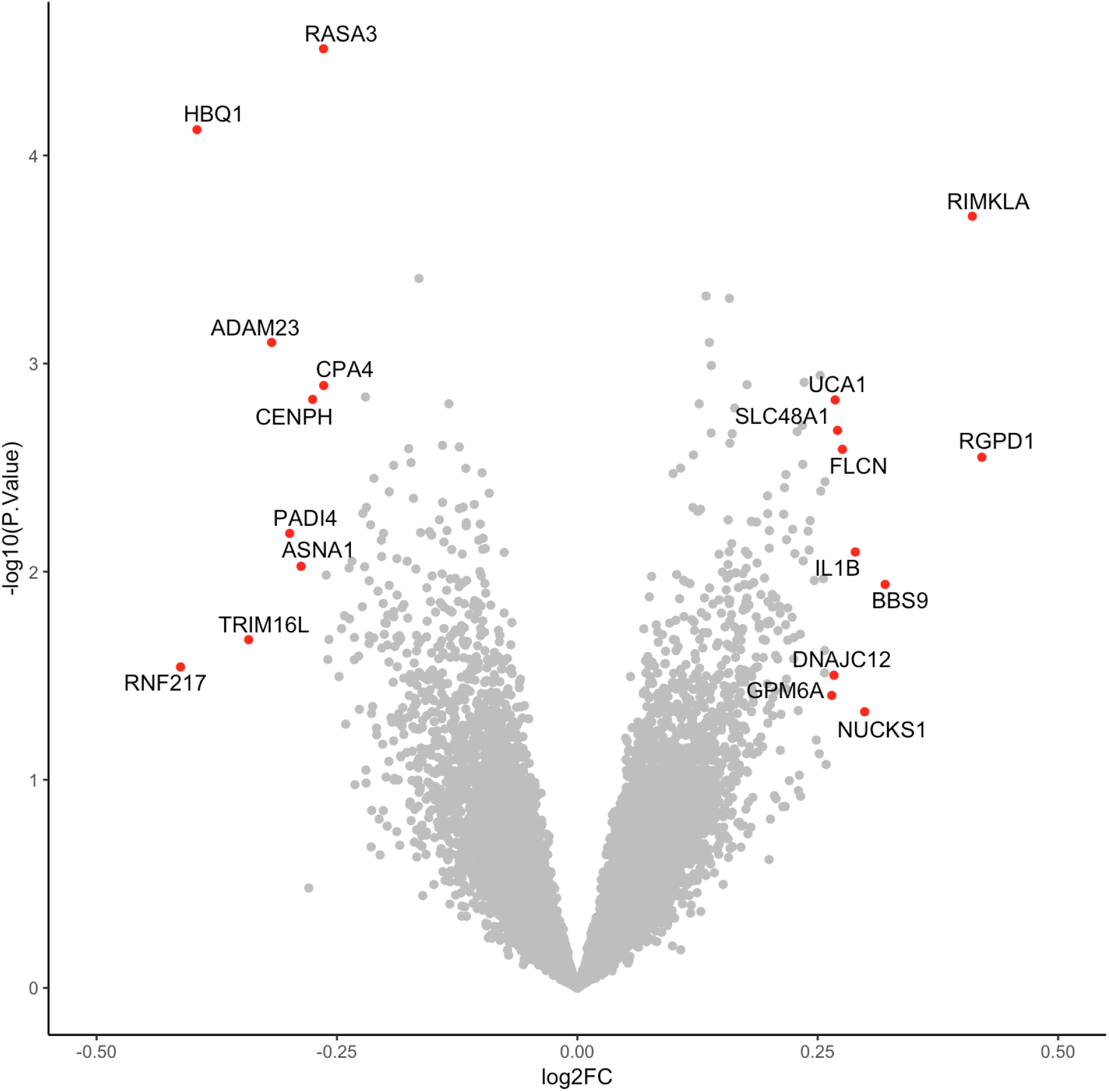
Differential transcriptional response to glucose among individuals with diabetes with and without retinopathy. Volcano plot summarizing genes exhibiting a differential response to glucose between individuals with diabetes with and without retinopathy (RG_PDR-nDR_). The difference in fold change between groups is represented on the X-axis and p-value of this difference on the Y-axis. Red indicates differential response genes (p-value < 0.05 and an absolute log_2_ FC difference > 0.26 (FC > 1.12)). FC fold change.

### Genes with differential response to glucose are implicated in the pathogenesis of diabetic retinopathy

We sought to assess whether the most significant differential response genes (RG_pdr-ndr_) could yield novel insights into diabetic retinopathy. An overview of our approach is presented in Figure 4a. First, we selected the top 103 genes (P < 0.01) that showed the largest difference in gene expression response to glucose between individuals with diabetes with and without retinopathy. We next identified all of the significant expression quantitative trait loci (eQTLs) for these genes in GTEx (version 7) [18]. In total, we found 7,253 unique eQTL SNPs (hereafter referred to as eSNPs) in at least one of the 48 tissues investigated by GTEx. Differential response genes are more likely to harbor eSNPs, and hence be eGenes, compared to the genome-wide average (P = 2.0 x 10^−16^) (Supplemental Figure S5). This suggests that differential response genes are more likely to be genetically regulated and may contribute to interindividual differences in the development of diabetic retinopathy. To test if the eSNPs for the 103 differential response genes were more associated with diabetic retinopathy than expected, we evaluated the association between the 7,253 differential response gene eSNPs and diabetic retinopathy using our published GWAS of diabetic retinopathy [8]. The 7,253 eSNPs from the differential response genes are enriched for association to diabetic retinopathy (FDR < 0.05) (Figure 4b). To further assess the significance of this enrichment, we performed permutation testing of eSNPs from random sets of 103 genes which demonstrated that less than 1% contained the same proportion of similarly skewed GWAS p-values (Supplemental Figure S6). The eSNPs for differential response genes were enriched among diabetic retinopathy meta-GWAS p-values relative to all eSNPs (P = 0.0012) and all SNPs (P = 0.0023) (Figure 4c). Thus, genes exhibiting a differential response to glucose (RG_pdr-ndr_) are associated with the development of severe diabetic retinopathy.

**Figure 4.**
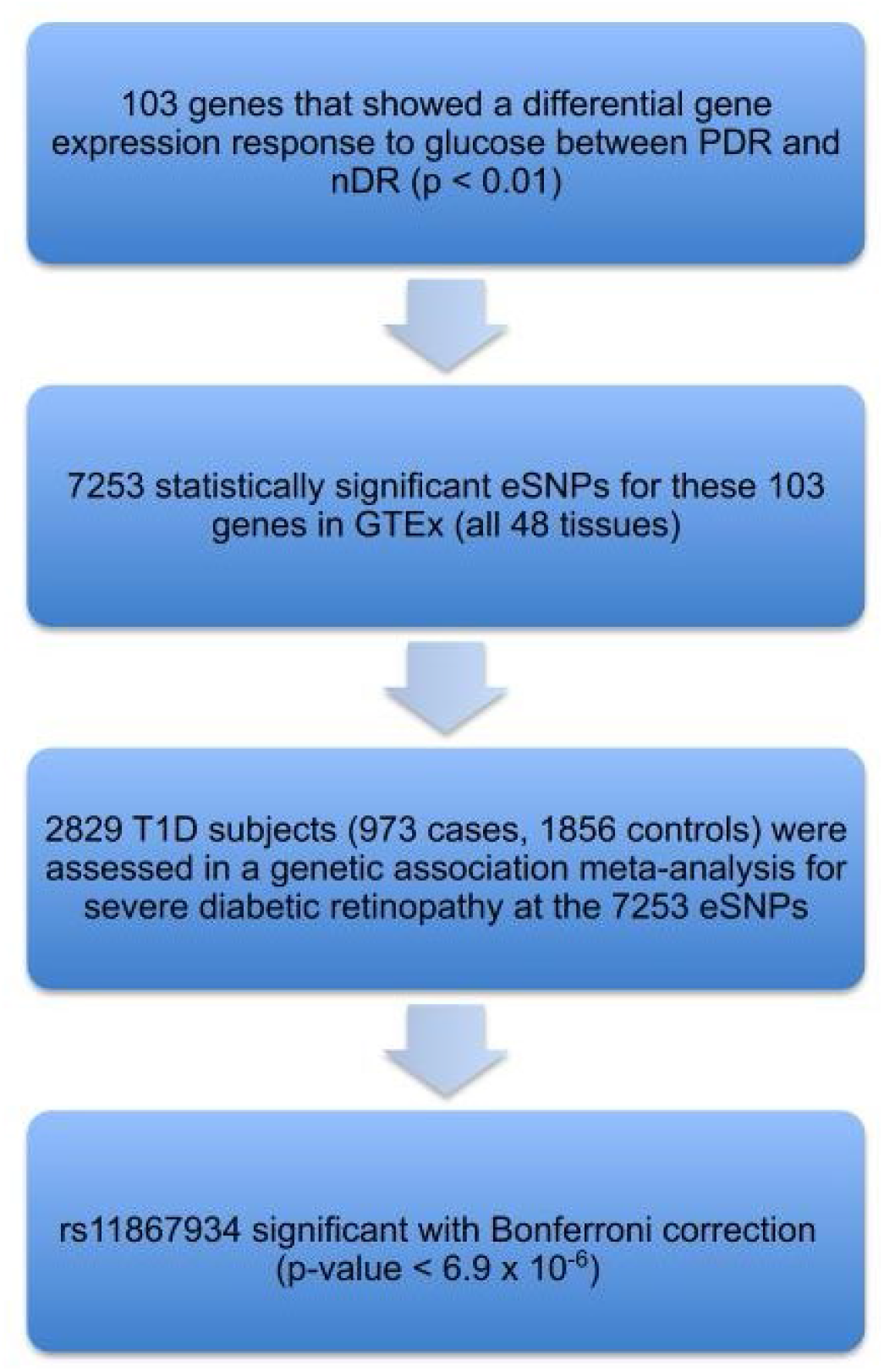
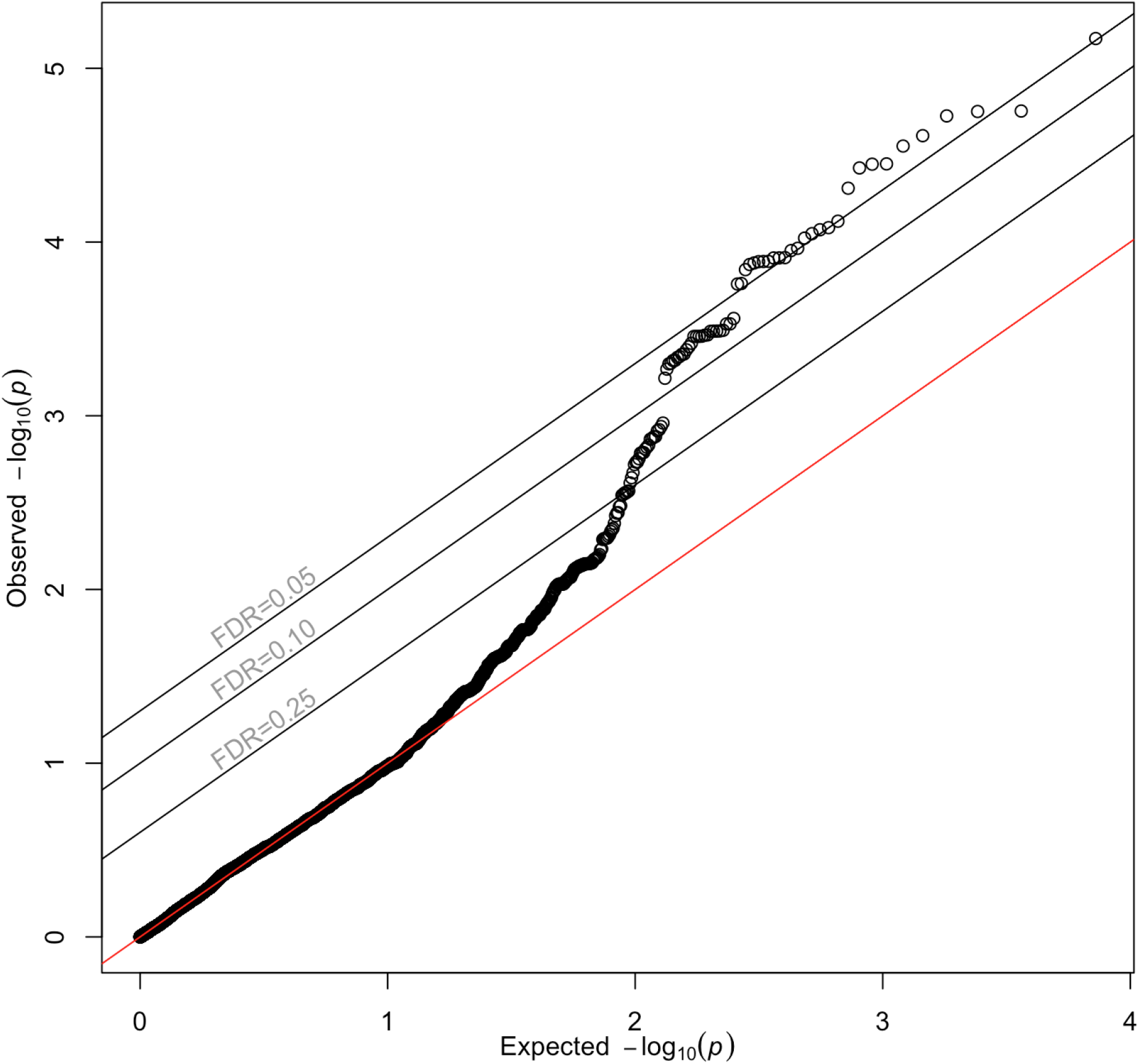
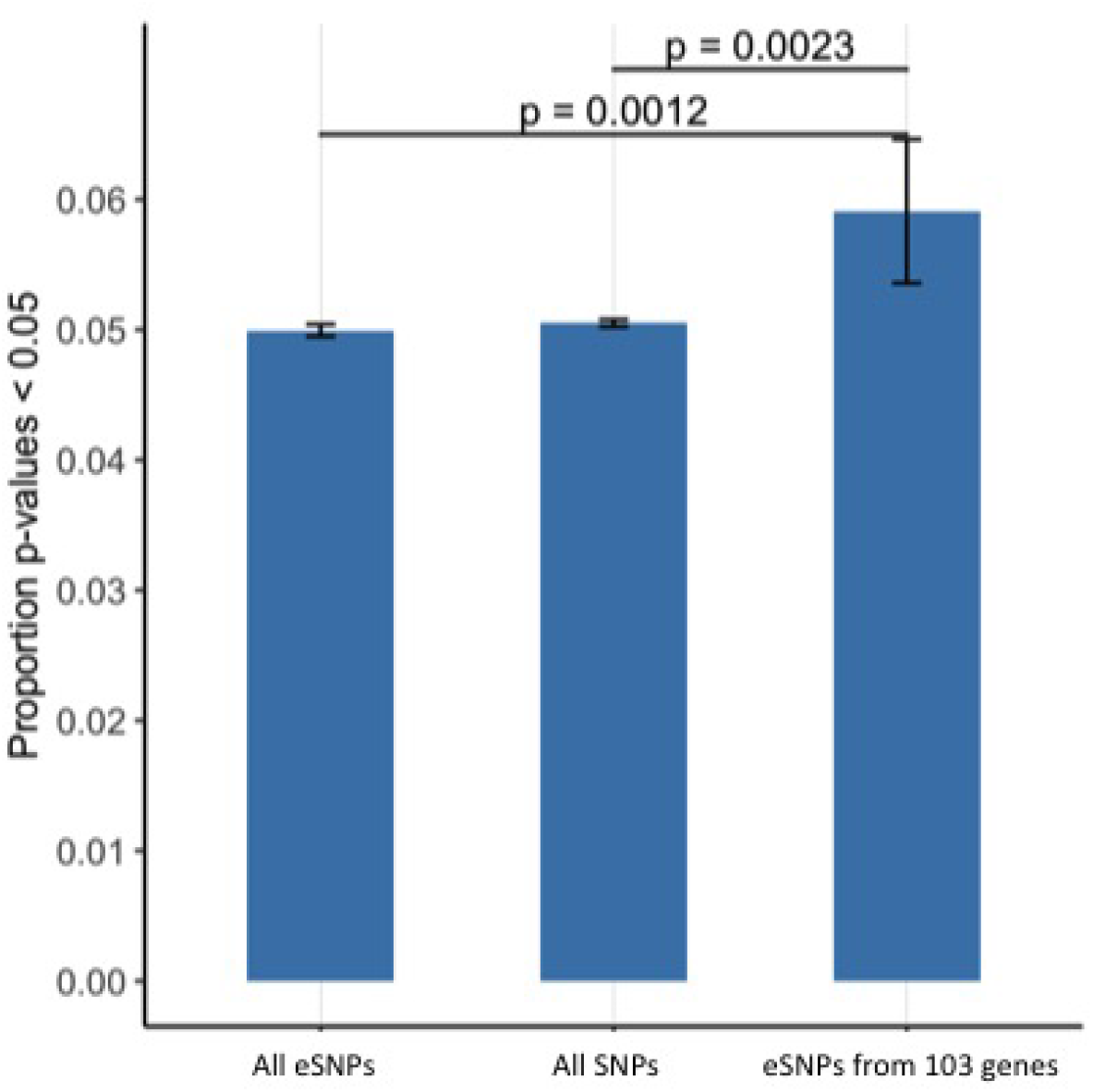
Association of glucose differential response genes (RG_pdr-ndr_) with diabetic retinopathy. a) Workflow of analytical steps integrating glucose differential response genes with genetic association to diabetic retinopathy. Flow chart showing key experimental steps based on stepwise findings. b) QQ Plot revealing a skew away from the null and above the FDR 0.05 threshold suggests that expression of some of the glucose response genes may be causally related to diabetic retinopathy. 7253 GTEx eSNPs were generated from the 103 differential response genes and tested for their association to diabetic retinopathy in a GWAS. Observed vs. Expected p-values are plotted. The null hypothesis of no difference between the observed and expected p-values is represented by the red line. No influence of population structure or other design factors was observed (genomic control inflation estimate □_GC_ = 1.005) [63]. c) Bar plot comparing frequency of p-values < 0.05 in diabetic retinopathy GWAS of: all eSNPs, all SNPs and eSNPs from the 103 differential response genes. An excess of GWAS p-values of < 0.05 is observed in the eSNPs from the glucose differential response genes (P = 0.0012 vs all eSNPs and P = 0.0023 vs all SNPs). The proportion of SNPs with P < 0.05 in the All SNPs, All eSNPs, and 103 differential response gene eSNPs are: 0.0505, 0.0499, and 0.0571 respectively.

### Folliculin (*FLCN*) is a putative diabetic retinopathy disease gene

The most significant retinopathy-associated eSNP, among the set of 7,253 eSNPs tested is rs11867934 (Figure 5a) FDR < 0.05; meta-GWAS P = 6.7×10^−6^ < Bonferroni adjusted p-value of 6.9×10^−6^; OR=0.86, 95%CI=0.71,1.00; Minor Allele Frequency, =0.22. rs11867934 is an intergenic eSNP for *FLCN* in multiple biologically relevant tissues including artery and nerve. We confirmed FLCN expression in the retina of human donor eyes (Supplemental Figure S7). In the LCLs derived from individuals with diabetes, *FLCN* was upregulated in response to glucose to a greater extent in individuals with diabetic retinopathy than in individuals with diabetes without retinopathy (log,FC difference = 0.276, P = 0.003) (Supplemental Figure S8).

**Figure 5.**
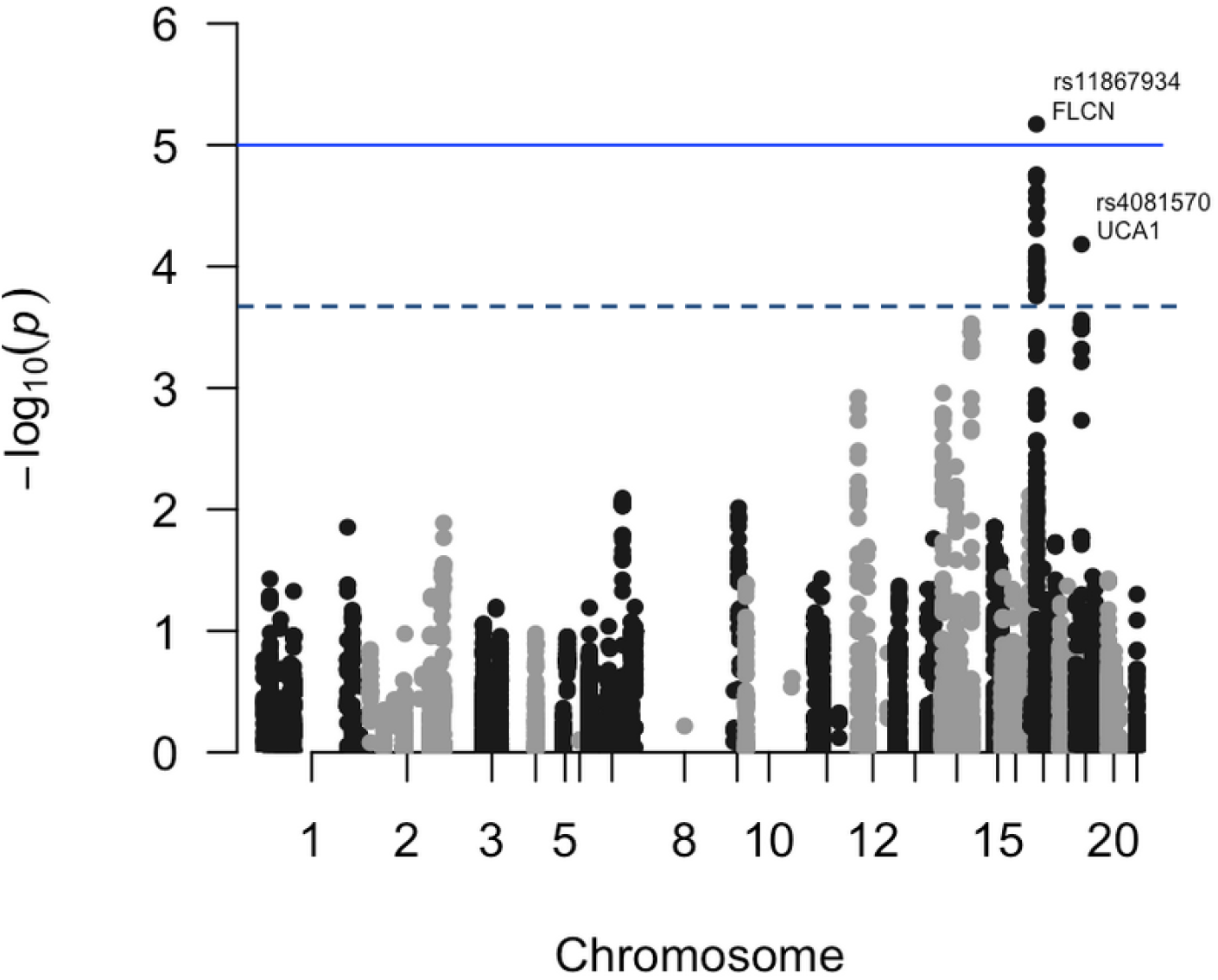
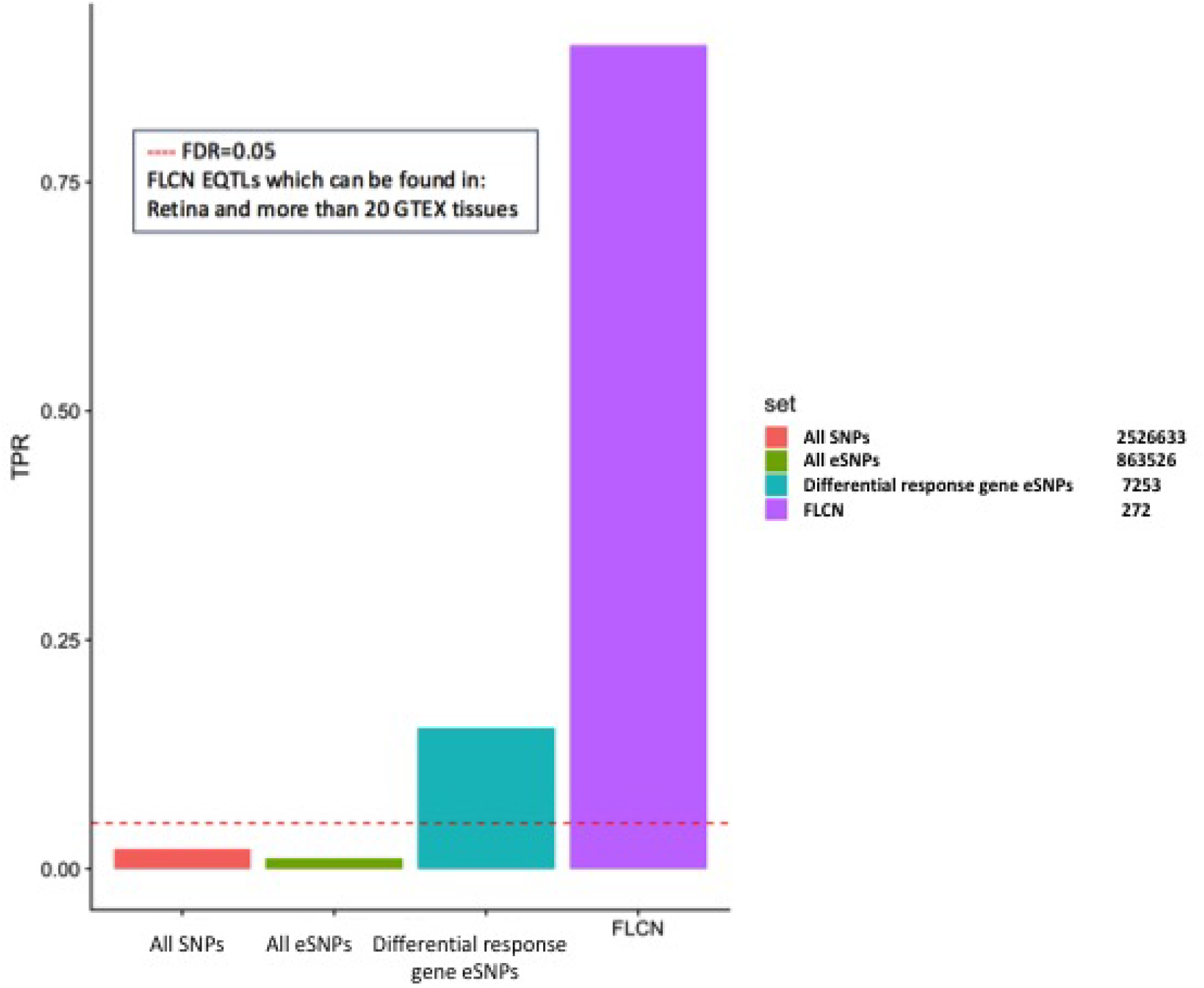
Diabetic retinopathy meta-GWAS for eSNPs of differential response genes to glucose. a) Manhattan plot of the results of the meta-GWAS for diabetic retinopathy showing association signals for the eSNPs from the differential response genes to glucose for individuals with and without retinopathy (RG_PDR-nDR_). Threshold lines represent Bonferroni correction (blue) and FDR < 0.05 (black). Association testing for diabetic retinopathy performed with 7253 eSNPs representing 103 differential response genes to glucose. b) Bar plot comparing the true positive rate (π_1_), TPR, for association of diabetic retinopathy with all SNPs, all eSNPs, eSNPs from the 103 differential response genes to glucose (n = 7,253), and eSNPs found in retina and > 20 GTEx tissues for folliculin (*FLCN*) (n = 272). TPR is an estimate of the proportion of tests that are true under the alternative hypothesis. Plot reveals significant enrichment for glucose response gene eSNPs in general and for *FLCN* eSNPs (π_1_ = 0.9) specifically.

eQTLs in retina have recently been mapped [30]. We determined that at least 43% of retina eQTLs are also eQTLs in GTEx LCLs. Examining the genome-wide association signal for a disease from eQTLs in aggregate can be a more powerful strategy to discern a heterogenous genetic signal than testing each of these SNPs individually. We collated all the eSNPs for *FLCN* in the retina. We assessed the aggregated association of *FLCN* eSNPs (N = 272 eSNPs significant in the retina and 20 or more GTEx tissues) to diabetic retinopathy in the meta-GWAS and observed an enrichment for association to diabetic retinopathy (π_1_ = 0.9; Figure 5b, Supplemental Figure S9). We then validated the *FLCN* association to diabetic retinopathy in a third cohort, the UK Biobank (UKBB) (Supplemental Table S3), and found that the *FLCN* eSNPs were enriched for association to diabetic retinopathy in the UKBB (π_1_ = 0.73) (Supplemental Figure S10).

We applied Mendelian randomization to assess whether the level of *FLCN* expression affects the development of diabetic retinopathy. We first imputed retinal *FLCN* expression in the UKBB, and then estimated the effects of the estimated *FLCN* expression on diabetic retinopathy using summary data-based Mendelian randomization analysis [36] (SMR). Mendelian randomization treats the genotype as an instrumental variable. A one standard deviation (SD) increase in the predicted retinal expression of *FLCN* increases the risk of diabetic retinopathy by 0.15 SD (95% confidence interval: 0.02 - 0.29, standard error 0.07, P = 0.024). Individuals with diabetes with high predicted retinal *FLCN* expression have increased odds of developing retinopathy (1.3 OR increase per SD increase in *FLCN* expression) [47]. We did not observe any evidence of horizontal pleiotropy (in which *FLCN* eSNPs are independently associated with both *FLCN* expression and diabetic retinopathy) confounding the analysis [HEIDI P > 0.05 (P = 0.2)] [36]. We detected an aggregated effect of 14 independent *FLCN* eQTLs (r^2^ < 0.2) on the development of diabetic retinopathy through *FLCN* expression using multi-SNP Mendelian randomization (P = 0.04) [37]. Together, these findings support the presence of genetic variation at the *FLCN* locus affecting both *FLCN* expression and the development of diabetic retinopathy through the expression of *FLCN*.

## Discussion

The cellular response to elevated glucose is an increasingly important pathway to understand in light of the emerging epidemic levels of diabetes worldwide [1]. Variations in the cellular response to glucose at a molecular level have not been well-characterized between cell types, and to an even lesser degree between individuals. In prior work, we characterized robust, repeatable interindividual differences in transcriptional response to glucose in LCLs of individuals with diabetic retinopathy [16]. As a lymphoblastoid cell line generated from each individual is genetically unique, it follows that the gene expression response to glucose between individuals should be phenotypically heterogeneous and that a portion of the interindividual variability will be genetically determined. We hypothesized that interindividual variation in the cellular response to glucose may reveal clues to the genetic basis of diabetic retinopathy, thereby providing insight into its predisposition.

We demonstrated that different individual-derived cell lines treated under identical culture conditions reveal an individual-specific transcriptional response to glucose and this signal far exceeds accompanying experimental noise. Transformation and multiple freeze/thaw passages do not homogenize the individualized response to high glucose induced gene expression in lymphoblastoid cell lines. Analyzing the individual glucose stimulated transcriptional response revealed several insights into the pathophysiology of the diabetic state and how it relates to the development of retinopathy. For instance, *TXNIP* was identified as the top differential response gene to glucose in all individuals (RG_all_). *TXNIP* is a key marker of oxidative stress. It is upregulated in the diabetic retina where it induces Muller cell activation [39]. High glucose treatment has been shown to increase *TXNIP* expression [40]. *TXNIP* is a glucose sensor whose expression has been strongly associated with both hyperglycemia and diabetic complications. Specifically, the *TXNIP* locus was differentially methylated in the primary leukocytes of EDIC cases and controls [40]. A key mechanism by which cells respond to stress is through changes in genome configuration. Conformational alterations in DNA packaging influence the accessibility of DNA for transcription. Structural changes in DNA conformation facilitate cellular adaptation and response to stimuli which can enable transcriptional changes. The gene set enrichment analysis showed that the cellular response to chronic glucose stress involves alterations in DNA accessibility which facilitates the gene expression response to this environmental stimulus [48]. The transcriptional response to glucose in part manifests as diminished immune responsiveness, a well characterized feature of diabetes [43] [49].

Further, we considered that the genetic component of an individual’s response to glucose may influence their susceptibility to diabetic complications like retinopathy. Cell lines from individuals with diabetes with and without retinopathy reveal differences in the response to glucose at a molecular level. In addition, not only were some of these differential response genes biologically relevant to diabetic retinopathy as exemplified by *IL1B* and *PDGF*, but also many had a genetic basis for their differential expression. By integrating the gene expression findings with GWAS data, we implicated folliculin (*FLCN*) as a putative disease gene in diabetic retinopathy. Mendelian randomization provided evidence that genetic variation affects diabetic retinopathy through alterations in *FLCN* expression thereby suggesting that FLCN expression is a mediator of diabetic retinopathy*. FLCN* is a biologically plausible diabetic retinopathy disease gene since its expression is present in both neuronal and vascular cells of the retina. Current evidence suggests that FLCN is a negative regulator of AMPK which helps to modulate the energy sensing ability of AMPK and plays a role in responding to cellular stress [50]. AMPK plays an important role in providing resistance to cellular stresses by regulating autophagy and cellular bioenergetics to avoid apoptosis. Loss of *FLCN* results in constitutive activation of AMPK. Higher levels of *FLCN* would suggest less cellular capacity to deal with stress [51]. Interestingly, the protective effect of agents such as metformin and fenofibrate on diabetic retinopathy might be mediated through AMPK [52] [53].

Our study design had several advantages over prior approaches aimed at revealing the genetic basis of diabetic retinopathy. First, we utilized white blood cells which are readily accessible from the peripheral circulation of human patients [48] and can reveal differential molecular characteristics depending on the stage of diabetic retinopathy [54] [55] [56]. Lymphoblastoid cell lines are derived from white blood cells making them a relevant cellular population to study for diabetic retinopathy. LCLs have been shown to be a powerful model system for functional genetic studies in humans [54, 56]. Second, a lymphoblastoid cell line (LCL) was generated for every individual enrolled in the landmark DCCT/EDIC study. DCCT/EDIC is the best-characterized prospective interventional cohort ever created to follow systemic complications of long-standing diabetes. DCCT/EDIC allows for detailed stratification of individuals, each of whom has had extensive prospective clinical phenotyping. Third, glucose was employed to elicit a provocative response in LCLs. By focusing on a secondary sequela of diabetes like retinopathy, the cellular response to glucose stimulation through transcription became a meaningful and directly relevant reflection of the stress each cell in the body encounters from diabetes. Insights into glucose stimulated gene expression in LCLs have broad applicability to multiple tissues of interest for diabetic complications (even in the retina as we have shown) due to significant evidence supporting a shared framework for gene regulation among tissues [18]. Finally, disease associated expression quantitative trait loci (eQTL) provide functional insights into the pathogenesis of a condition. We show that altering the levels of *FLCN* expression impacts risk of diabetic retinopathy. Aggregating independent eQTLs for the same gene (that are not in high linkage disequilibrium) revealed an enriched association that may otherwise have been missed by a conventional GWAS approach [57]. Treating the associated eQTL as an instrumental variable, Mendelian randomization supported the causality of *FLCN* in the pathogenesis of the disease. Inherently, this approach yielded all three M’s of target modulation: mechanism, magnitude and markers [58].

The present work had inherent limitations. First, LCLs are not primary cells but rather a transformed cell line. The Multiple Tissue Human Expression Resource (MuTHER) LCL study revealed a large impact of common environmental exposure, stemming from shared sample handling, on gene expression in twin LCLs [59]. The significant correlation of these extrinsic factors on LCL gene expression emphasizes the importance of randomization and technical replicates which we implemented in this study. Moreover, as a cell line, heterogeneous genomic alterations have been identified in lymphoblastoid cells that increase with passaging, thereby raising the concern that this can lead to variability in their transcriptome [60]. Importantly, the EDIC cell lines employed in this study were only passaged once previously. Additionally, genomic changes have only a minor effect on genotypic frequencies with a 99.63% genotype concordance between lymphoblastoid cells and their parent leukocytes. Mendelian error rates in levels of heterozygosity are not significantly different between lymphoblastoid cell lines and their primary B-lymphocyte cells of origin [61]. Second, it is not possible to delineate cause from effect in gene expression studies. Gene expression changes may be causal, epiphenomena, or due to reverse causality (the disease causing the gene expression changes rather than the other way around). In this study, by integrating genetic analyses with gene expression and recognizing that variation in the underlying genome precedes disease onset and can therefore be considered an instrumental variable, we identified through Mendelian randomization potentially causal gene expression changes in *FLCN* that act as a mediator for retinopathy thereby avoiding the trap of reverse causality. Finally, eQTL found in LCLs may not be relevant to diabetic retinopathy. As noted previously we found 43% of retina eQTL are shared with LCLs. We demonstrated that independent *FLCN* eQTLs found both in the retina and GTEx tissues showed an enriched association to diabetic retinopathy, a finding that was replicated in a large independent cohort from the UK Biobank. For complex trait associations in general and for those specifically in the retina, eQTL that are shared between tissues explain a greater proportion of associations than tissue specific eQTL [15]. For instance, shared tissue eQTL are enriched among genetic associations to age-related macular degeneration, another common retinal disease, despite the high tissue specificity of the disease [30] [62].

In summary, integration of gene expression from a relevant cellular model with genetic association data provided insights into the functional relevance of genetic risk for a complex disease. Using disease associated differential gene and eQTL based genomewide association testing, we identified causal genetic pathways for diabetic retinopathy. Specifically, our studies implicated *FLCN* as a putative diabetic retinopathy susceptibility gene. Future work that incorporates more extensive molecular profiling of the cellular response to glucose in conjunction with a greater number of cell lines may yield further insights into the underlying genetic basis of diabetic retinopathy.

## Acknowledgements

This research has been conducted using the UK Biobank Resource under Application Number 44316.

We acknowledge the guidance and assistance provided by the members of the DCCT/EDIC committee at the time of this publication. A complete list of investigators and members of the Research Group appears in *N Engl J Med* 2017, 376:1507-1516.

We thank the DNA Services Facility and the Research Histology and Tissue Imaging Core at UIC Research Resources Center for assistance in histological techniques and image acquisition.

We thank Andrew D. Paterson, MD for helpful input and comments.

## References

1. National Diabetes Fact Sheet. Centers for Disease Control and Prevention [webpage] 2011 11-02-2011]; Available from: http://www.cdc.gov/diabetes/pubs/estimates11.htm#12.

2. https://www.cdc.gov/features/diabetic-retinopathy/index.html.

3. Group, D.E.R., et al., Frequency of Evidence-Based Screening for Retinopathy in Type 1 Diabetes. N Engl J Med, 2017. 376(16): p. 1507–1516.

4. Sun, J.K., et al., Protection from retinopathy and other complications in patients with type 1 diabetes of extreme duration: the joslin 50-year medalist study. Diabetes Care, 2011. 34(4): p. 968–74.

5. Gao, X., et al., Native American ancestry is associated with severe diabetic retinopathy in Latinos. Invest Ophthalmol Vis Sci, 2014. 55(9): p. 6041–5.

6. Arar, N.H., et al., Heritability of the severity of diabetic retinopathy: the FIND-Eye study. Invest Ophthalmol Vis Sci, 2008. 49(9): p. 3839–45.

7. Hietala, K., et al., Heritability of proliferative diabetic retinopathy. Diabetes, 2008. 57(8): p. 2176–80.

8. Grassi, M.A., et al., Genome-wide meta-analysis for severe diabetic retinopathy. Human molecular genetics, 2011. 20(12): p. 2472–81.

9. Grassi, M.A., et al., Replication analysis for severe diabetic retinopathy. Investigative ophthalmology & visual science, 2012. 53(4): p. 2377–81.

10. Pollack, S., et al., Multiethnic Genome-Wide Association Study of Diabetic Retinopathy Using Liability Threshold Modeling of Duration of Diabetes and Glycemic Control. Diabetes, 2019. 68(2): p. 441–456.

11. Fritsche, L.G., et al., Age-related macular degeneration: genetics and biology coming together. Annu Rev Genomics Hum Genet, 2014. 15: p. 151–71.

12. Fritsche, L.G., et al., A large genome-wide association study of age-related macular degeneration highlights contributions of rare and common variants. Nat Genet, 2016. 48(2): p. 134–43.

13. Risch, N. and K. Merikangas, The future of genetic studies of complex human diseases. Science, 1996. 273(5281): p. 1516–7.

14. Maurano, M.T., et al., Systematic localization of common disease-associated variation in regulatory DNA. Science, 2012. 337(6099): p. 1190–5.

15. Gamazon, E.R., et al., Using an atlas of gene regulation across 44 human tissues to inform complex disease- and trait-associated variation. Nat Genet, 2018. 50(7): p. 956–967.

16. Grassi, M.A., et al., Lymphoblastoid Cell Lines as a Tool to Study Inter-Individual Differences in the Response to Glucose. PLoS One, 2016. 11(8): p. e0160504.

17. Grassi, M.A., et al., Genetic Variation Is the Major Determinant of Individual Differences in Leukocyte Endothelial Adhesion. PLoS ONE, 2014. 9(2): p. e87883.

18. Consortium, G., Human genomics, The Genotype-Tissue Expression (GTEx) pilot analysis: multitissue gene regulation in humans. Science, 2015. 348(6235): p. 648–60.

19. Effect of intensive therapy on the microvascular complications of type 1 diabetes mellitus. JAMA : the journal of the American Medical Association, 2002. 287(19): p. 2563–9.

20. Epidemiology of Diabetes Interventions and Complications (EDIC). Design, implementation, and preliminary results of a long-term follow-up of the Diabetes Control and Complications Trial cohort. Diabetes care, 1999. 22(1): p. 99–111.

21. (DCCT)., T.D.C.a.C.T., The Diabetes Control and Complications Trial (DCCT). Design and methodologic considerations for the feasibility phase. The DCCT Research Group. Diabetes, 1986. 35(5): p. 530–45.

22. The effect of intensive diabetes treatment on the progression of diabetic retinopathy in insulin-dependent diabetes mellitus. The Diabetes Control and Complications Trial. Archives of ophthalmology, 1995. 113(1): p. 36–51.

23. Caramori, M.L., et al., Gene expression differences in skin fibroblasts in identical twins discordant for type 1 diabetes. Diabetes, 2012. 61(3): p. 739–44.

24. Du, P., W.A. Kibbe, and S.M. Lin, lumi: a pipeline for processing Illumina microarray. Bioinformatics, 2008. 24(13): p. 1547–8.

25. Lin, S.M., et al., Model-based variance-stabilizing transformation for Illumina microarray data. Nucleic Acids Res, 2008. 36(2): p. e11.

26. Choy, E., et al., Genetic analysis of human traits in vitro: drug response and gene expression in lymphoblastoid cell lines. PLoS genetics, 2008. 4(11): p. e1000287.

27. Ritchie, M.E., et al., limma powers differential expression analyses for RNA-sequencing and microarray studies. Nucleic Acids Res, 2015. 43(7): p. e47.

28. Becker, R.A., Chambers, J. M. and Wilks, A. R., The New S Language. Wadsworth & Brooks/Cole, 1988.

29. Subramanian, A., et al., Gene set enrichment analysis: a knowledge-based approach for interpreting genome-wide expression profiles. Proc Natl Acad Sci U S A, 2005. 102(43): p. 15545–50.

30. Ratnapriya, R., et al., Retinal transcriptome and eQTL analyses identify genes associated with age-related macular degeneration. Nat Genet, 2019.

31. Grassi, M.A., et al., Validity of Self-Report in Type 1 Diabetic Subjects for Laser Treatment of Retinopathy. Ophthalmology, 2013.

32. Sudlow, C., et al., UK biobank: an open access resource for identifying the causes of a wide range of complex diseases of middle and old age. PLoS Med, 2015. 12(3): p. e1001779.

33. Grassi, M.A., et al., Patient self-report of prior laser treatment reliably indicates presence of severe diabetic retinopathy. American journal of ophthalmology, 2009. 147(3): p. 501–4.

34. Chang, C.C., et al., Second-generation PLINK: rising to the challenge of larger and richer datasets. Gigascience, 2015. 4: p. 7.

35. Davies, N.M., et al., Multivariable two-sample Mendelian randomization estimates of the effects of intelligence and education on health. Elife, 2019. 8.

36. Zhu, Z., et al., Integration of summary data from GWAS and eQTL studies predicts complex trait gene targets. Nat Genet, 2016. 48(5): p. 481–7.

37. Wu, Y., et al., Integrative analysis of omics summary data reveals putative mechanisms underlying complex traits. Nat Commun, 2018. 9(1): p. 918.

38. Storey, J.D. and R. Tibshirani, Statistical significance for genomewide studies. Proc Natl Acad Sci U S A, 2003. 100(16): p. 9440–5.

39. Devi, T.S., et al., TXNIP regulates mitophagy in retinal Muller cells under high-glucose conditions: implications for diabetic retinopathy. Cell Death Dis, 2017. 8(5): p. e2777.

40. Chen, Z., et al., Epigenomic profiling reveals an association between persistence of DNA methylation and metabolic memory in the DCCT/EDIC type 1 diabetes cohort. Proc Natl Acad Sci U S A, 2016. 113(21): p. E3002–11.

41. Delamaire, M., et al., Impaired leucocyte functions in diabetic patients. Diabet Med, 1997. 14(1): p. 29–34.

42. Al-Mashat HA, K.S., Liu R, Behl Y, Desta T, Graves DT, Diabetes enhances mRNA levels of proapoptotic genes and caspase activity, which contribute to impaired healing. Diabetes., 2006. 55(2): p. 487–95.

43. Shanmugam, N., et al., High glucose-induced expression of proinflammatory cytokine and chemokine genes in monocytic cells. Diabetes, 2003. 52(5): p. 1256–64.

44. Liu, Y., M. Biarnes Costa, and C. Gerhardinger, IL-1beta Is Upregulated in the Diabetic Retina and Retinal Vessels: Cell-Specific Effect of High Glucose and IL-1beta Autostimulation. PloS one, 2012. 7(5): p. e36949.

45. Freyberger H, B.M., Yakut H, Hammer J, Effert R, Schifferdecker E, Schatz and D.M. H, Increased levels of platelet-derived growth factor in vitreous fluid of patients with proliferative diabetic retinopathy. Exp Clin Endocrinol Diabetes, 2000. 108(2):106–9.

46. Hammes, H.P., et al., Pericytes and the pathogenesis of diabetic retinopathy. Diabetes, 2002. 51(10): p. 3107–12.

47. S., C., A simple method for converting an odds ratio to effect size for use in meta-analysis. Stat Med., 2000. Nov 30;19(22):3127–31.

48. Smith, K.T. and J.L. Workman, Chromatin proteins: key responders to stress. PLoS Biol, 2012. 10(7): p. e1001371.

49. Mowat, A. and J. Baum, Chemotaxis of polymorphonuclear leukocytes from patients with diabetes mellitus. N Engl J Med, 1971. 284(12): p. 621–7.

50. Hasumi, H., et al., Regulation of mitochondrial oxidative metabolism by tumor suppressor FLCN. J Natl Cancer Inst, 2012. 104(22): p. 1750–64.

51. Possik, E., et al., Folliculin regulates ampk-dependent autophagy and metabolic stress survival. PLoS Genet, 2014. 10(4): p. e1004273.

52. Kim J, A.J., Kim JH, Yu YS, Kim HS, Ha J, Shinn SH, Oh YS. 2007 Epub 2007 Jan 27., Fenofibrate regulates retinal endothelial cell survival through the AMPK signal transduction pathway. Exp Eye Res, 2007. May;84(5):886–93.

53. Joe, S.G., et al., Anti-angiogenic effect of metformin in mouse oxygen-induced retinopathy is mediated by reducing levels of the vascular endothelial growth factor receptor Flk-1. PLoS One, 2015. 10(3): p. e0119708.

54. Tang, J. and T.S. Kern, Inflammation in diabetic retinopathy. Progress in retinal and eye research, 2011.

55. Gubitosi-Klug, R.A., et al., 5-Lipoxygenase, but not 12/15-lipoxygenase, contributes to degeneration of retinal capillaries in a mouse model of diabetic retinopathy. Diabetes, 2008. 57(5): p. 1387–93.

56. Kern, T.S., Contributions of inflammatory processes to the development of the early stages of diabetic retinopathy. Experimental diabetes research, 2007. 2007: p. 95103.

57. Wu, L., et al., A transcriptome-wide association study of 229,000 women identifies new candidate susceptibility genes for breast cancer. Nat Genet, 2018. 50(7): p. 968–978.

58. Plenge, R.M., Priority index for human genetics and drug discovery. Nat Genet, 2019. 51(7): p. 1073–1075.

59. Wright, F.A., et al., Heritability and genomics of gene expression in peripheral blood. Nat Genet, 2014. 46(5): p. 430–7.

60. Ben-David, U., et al., Genetic and transcriptional evolution alters cancer cell line drug response. Nature, 2018. 560(7718): p. 325–330.

61. McCarthy, N.S., et al., Integrity of genome-wide genotype data from low passage lymphoblastoid cell lines. Genom Data, 2016. 9: p. 18–21.

62. Unlu, G., et al., GRIK5 Genetically Regulated Expression Associated with Eye and Vascular Phenomes: Discovery through Iteration among Biobanks, Electronic Health Records, and Zebrafish. Am J Hum Genet, 2019. 104(3): p. 503–519.

63. Devlin B, R.K., Genomic control for association studies. Biometrics. 1999;55:997–1004.

