## Supplemental information for "Mendelian randomization identifies folliculin expression as a mediator of diabetic retinopathy"

**Tables:**

**Table S1. Demographic features of the DCCT/EDIC study subjects with type 1 diabetes**

| | nDR | PDR |
|---|---|---|
| **N** | **7** | **8** |
| **Mean age in years at DCCT baseline* (yrs, std)** | **32 (3.9)** | **31 (8.5)** |
| **Caucasian Ethnicity (%)** | **100** | **100** |
| **Female (%)** | **4 (57.1)** | **5 (62.5)** |
| **Mean duration of type 1 diabetes in months at DCCT baseline (yrs, std)** | **27 (13.4)** | **53 (43.4)** |
| **HbA1c$^{+}$ (%, std)** | **7.62 (1.07)** | **9.71 (2.37)** |
| **Intensive Treatment group^ (%)** | **2 (28.6)** | **3 (37.5)** |
| **Secondary Intervention Cohort# (%)** | **0 (0)** | **1 (12.5)** |

All subjects from DCCT/EDIC – the Diabetes Control and Complications Trial/ Epidemiology of Diabetes Interventions and Complications cohort
nDR - no Diabetic Retinopathy; PDR - proliferative diabetic retinopathy
* DCCT baseline at subject enrollment (1983–1989)
(std) standard deviation
$^{+}$HbA1c mean includes all DCCT visits except the baseline visit.
^ For the duration of the DCCT study the intensive treatment group maintained a HbA1c of approximately 7% as compared to approximately 9% in the conventional treatment group.
#The Secondary Intervention Cohort consisted of subjects with type 1 diabetes for 1-15 years and mild to moderate non-proliferative retinopathy and a urinary albumin excretion rate < 200 mg/dl at baseline. The Primary Prevention cohort consisted of subjects with type 1 diabetes for 1-5 years and no diabetes related complications.

**Table S2. Demographic features of individuals without diabetes from the Coriell Institute for Medical Research NIGMS Human Genetic Cell Repository**

| Subject | Ethnicity | Gender | Age*(years) | BMI | Notes |
|---|---|---|---|---|---|
| GM14581 | Caucasian | Male | 18 | 24 | |
| GM14569 | Caucasian | Male | 24 | N/A | |
| GM14381 | Caucasian | Female | 20 | 21 | # |
| GM07012 | Caucasian | Female | N/A | N/A | CEPH[%] |
| GM14520 | Caucasian | Female | 22 | 32 | |
| GM11985 | Caucasian | Female | N/A | N/A | CEPH[%] |
| GM07344 | Caucasian | Female | N/A | N/A | CEPH[%] |

Notes:
* At time of sampling
### Family history of diabetes
% Repository Linkage Families
N/A Not Available

**Table S3. Demographic features of UK Biobank subjects with diabetes used in the diabetic retinopathy GWAS**

| | Cases (n=2,332) | Controls (n=14,680) |
|---|---|---|
| Age years | 60 (6.86) | 60 (7.01) |
| HbA1c % | 7.4 (3.5) | 6.8 (3.4) |
| Female % | 34.70 | 38.50 |
| T1D (n) | 8% (187) | 3.30% (484) |
| T2D (n) | 76.70% (1789) | 67.90% (9968) |
| Unspecified (n) | 15.3% (356) | 28.8% (4228) |

Notes:
Age and HbA1c at enrollment given with mean and (standard deviation).
T1D: Type 1 diabetes (data defined as coded in ICD10 as E10)
T2D: Type 2 diabetes (data defined as coded in ICD10 as E11)
Duration of diabetes is not available.
Case subjects were defined as those who answered “yes” to questionnaire data eyesight field 6148 ‘Diabetes related eye disease’. Control subjects were defined as those who answered “yes” to data field 2443 ‘Diabetes diagnosed by doctor, excluding case subjects.

**Table S4. Differential response to Glucose PDR vs nDR ($RG_{pdr-ndr}$)**

| geneSymbol | Gene Name | Log2FC difference | p-value |
|---|---|---|---|
| RASA3 | RAS p21 protein activator 3 | -0.27 | $3.1x10^{-5}$ |
| HBQ1 | hemoglobin subunit theta 1 | -0.40 | $7.6x10^{-5}$ |
| RIMKLA | ribosomal modification protein rimK like family member A | 0.42 | $2x10^{-4}$ |
| ADAM23 | ADAM metallopeptidase domain 23 | -0.32 | $8x10^{-4}$ |
| CPA4 | carboxypeptidase A4 | -0.27 | $1.3x10^{-3}$ |
| CENPH | centromere protein H | -0.28 | $1.5x10^{-3}$ |
| UCA1 | urothelial cancer associated 1 | 0.27 | $1.5x10^{-3}$ |
| SLC48A1 | solute carrier family 48 member 1 | 0.28 | $2.1x10^{-3}$ |
| FLCN | folliculin | 0.28 | $2.6x10^{-3}$ |
| RGPD1 | RANBP2-like and GRIP domain containing 1 | 0.43 | $2.9x10^{-3}$ |
| PADI4 | peptidyl arginine deiminase 4 | -0.30 | $6.6 x10^{-3}$ |
| IL1B | interleukin 1 beta | 0.29 | $8.1x10^{-3}$ |
| ASNA1 | arsA (bacterial) arsenite transporter, ATP-binding, homolog 1 | -0.29 | $9.5x10^{-3}$ |
| BBS9 | Bardet-Biedl syndrome 9 | 0.33 | $1.2x10^{-2}$ |
| TRIM16L | tripartite motif containing 16-like | -0.35 | $2.2x10^{-2}$ |
| RNF217 | ring finger protein 217 | -0.42 | $2.9x10^{-2}$ |
| DNAJC12 | DnaJ heat shock protein family (Hsp40) member C12 | 0.27 | $3.2x10^{-2}$ |
| GPM6A | glycoprotein M6A | 0.27 | $4x10^{-2}$ |
| NUCKS1 | nuclear casein kinase and cyclin dependent kinase substrate 1 | 0.30 | $4.8x10^{-2}$ |

*List includes those genes with an absolute $log_2$ FC > 0.26 difference between the two groups and an uncorrected p-value of < 0.05. PDR – subject with proliferative diabetic retinopathy. nDR – subject with diabetes but no retinopathy.

**Figures:**

**Fig. S1. P-value distribution for transcriptional response to glucose in all 22 individuals ($RG_{All}$) (No diabetes, nDR and PDR).**

Plotted are limma-derived differential expression p-values for 11,548 genes. The dashed line represents the expected null distribution.

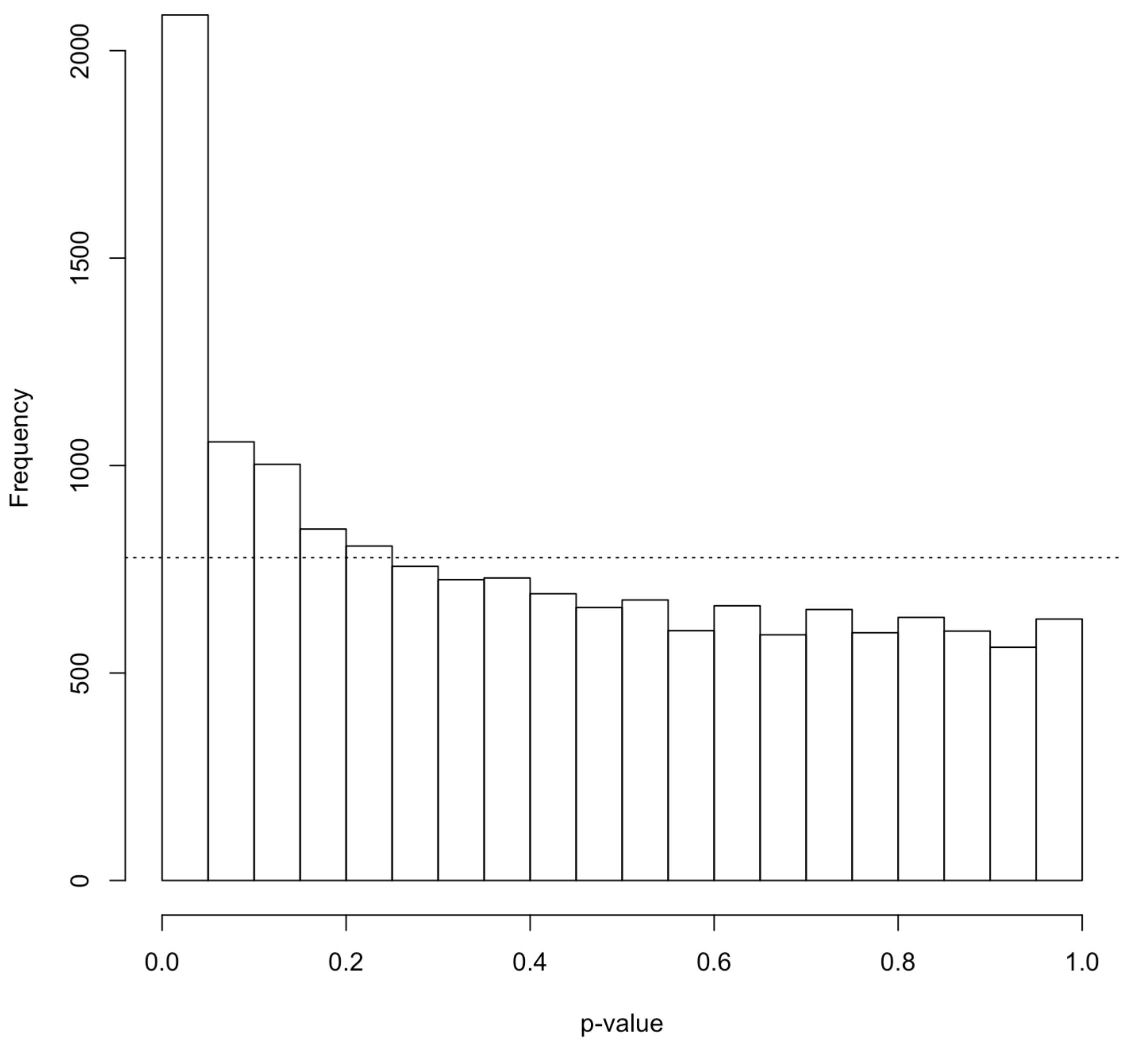

**Fig. S2. Intra- and interindividual transcriptome variation in high glucose treatment.**

Intraindividual variation in the transcriptome was quantified among biological replicates and compared to interindividual variation, for individual genes (A,B), and comparing the two distributions of all genes (B,C) .

**a)**

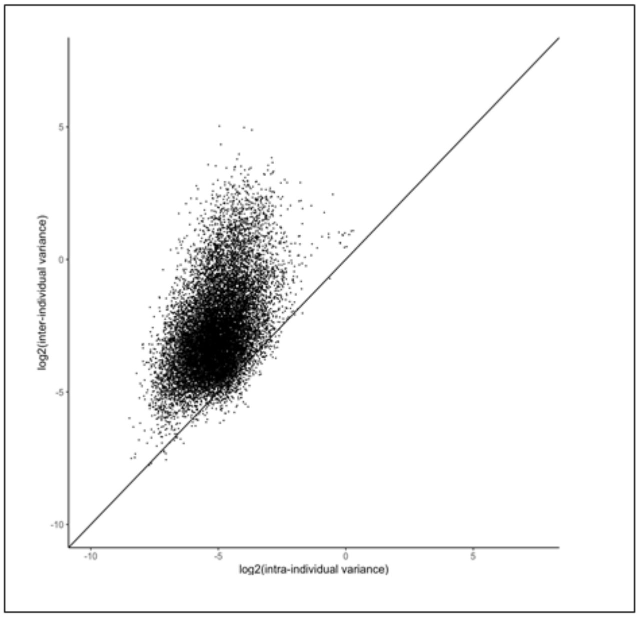


**b)**

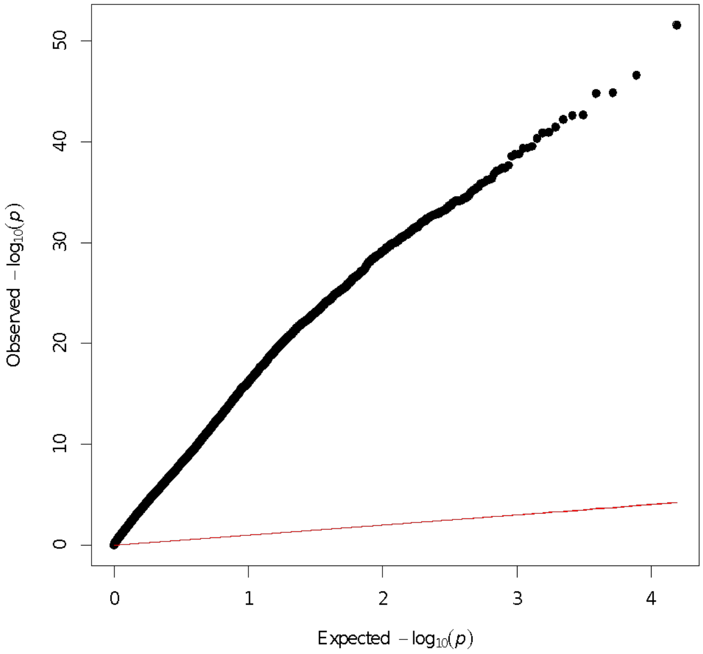

c)

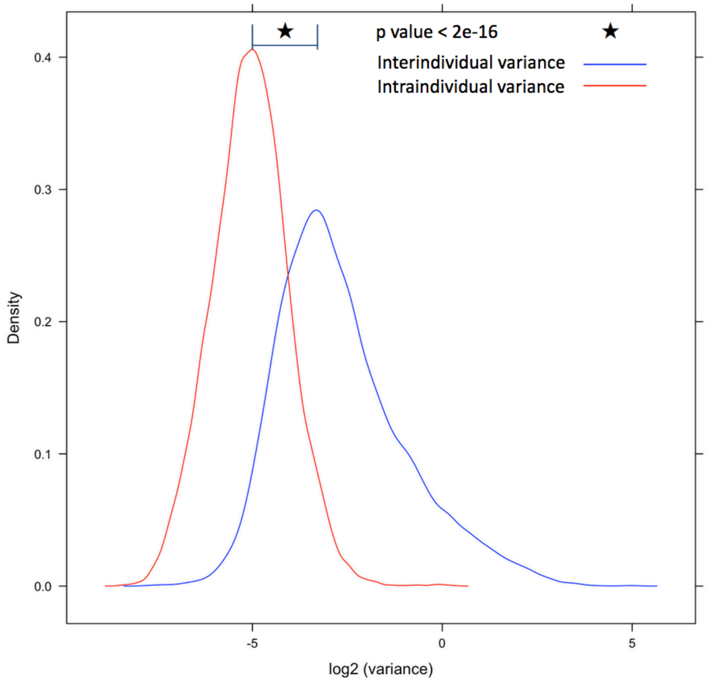
p value < 2e-16
Interindividual variance
Intraindividual variance
Density
0.4
0.3
0.2
0.1
0.0
-5
0
5
log2 (variance)

**Fig. S3. Multidimensional scaling based on differential response to glucose (rg).**

Each point represents a single study subject; individuals with proliferative diabetic retinopathy (PDR, red, n=8) and individuals with diabetes without retinopathy (nDR, blue, n=7). The first coordinate (Dim.1, x-axis) is correlated with subject retinopathy status ($RG_{pdr-ndr}$, P = 3x $10^{-6}$).

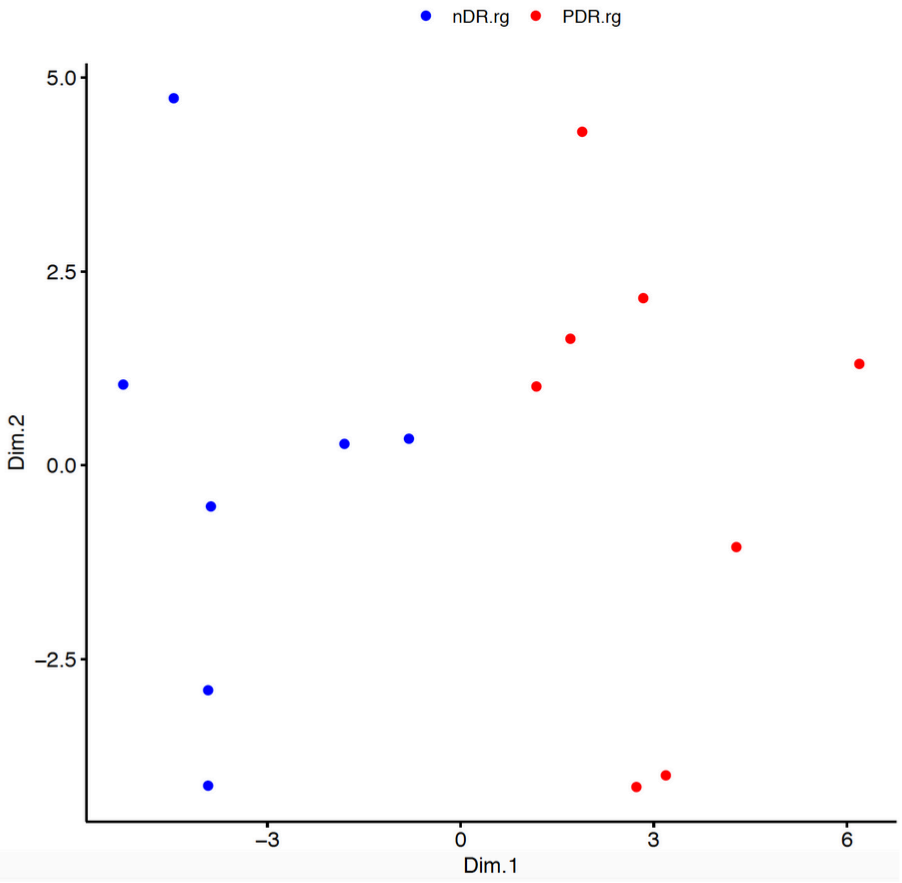

**Fig. S4. Gene Set Enrichment Analysis (GSEA) of genes with differential response to glucose between individuals with diabetes with and without diabetic retinopathy.**

Normalized enrichment score (NES)

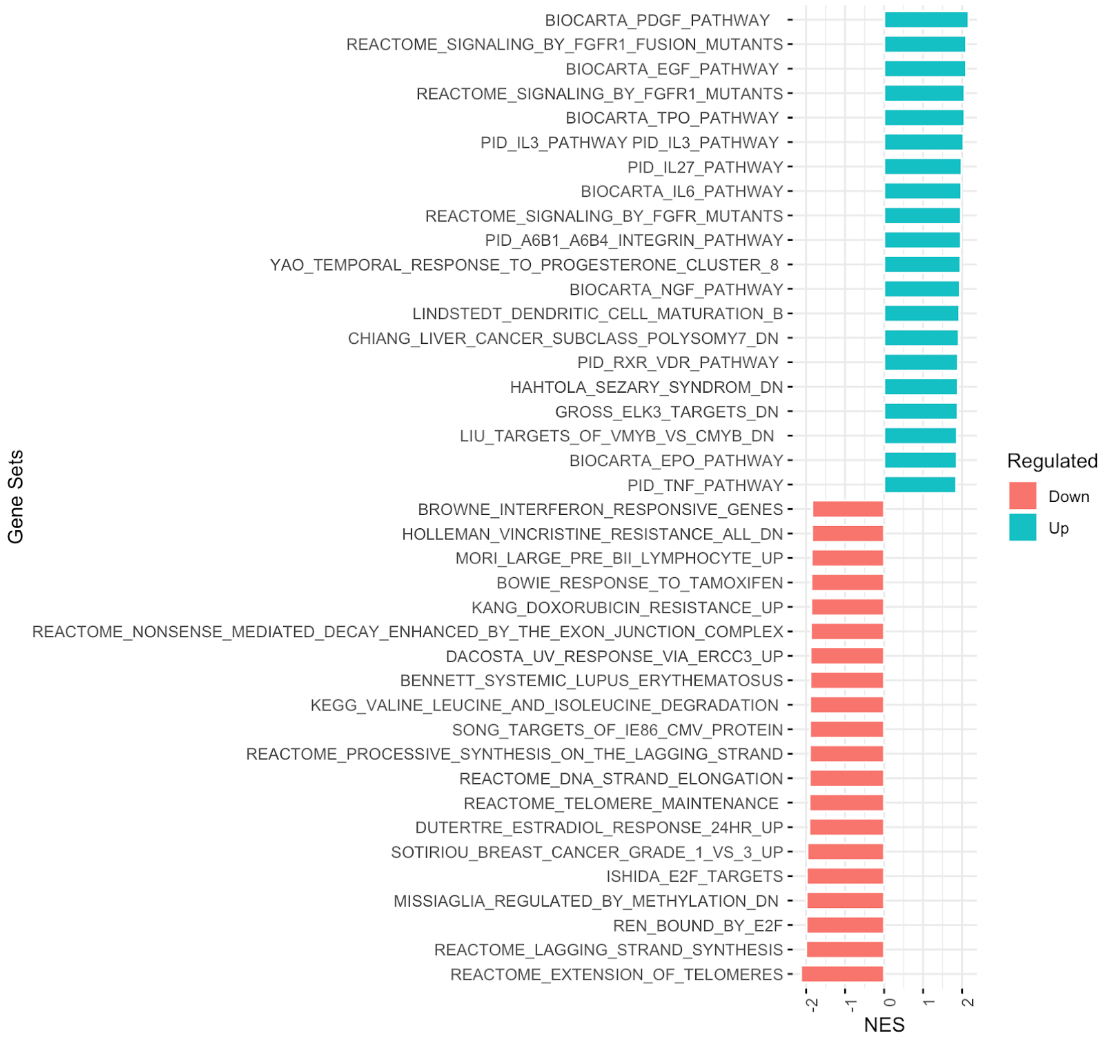

**Fig. S5. Enrichment of eGenes in glucose differential response genes.**

The proportion of eGenes is plotted to compare all 12,503 genes assessed on the microarray to the 103 glucose response genes. An eGene is defined as any gene with a GTEx eSNP (q-value < 0.05) in any tissue. GTEx version 7.

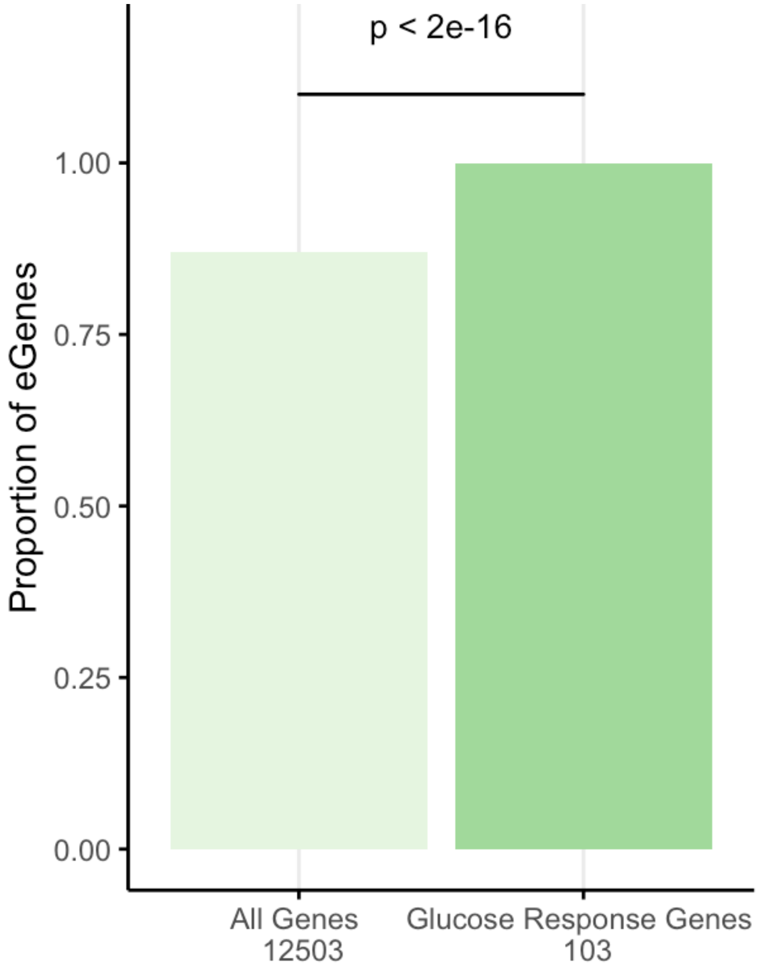

**Fig S6: Histogram of frequency in which permutations of eSNPs generated from random sets of 103 genes revealed similar p-values to those generated from the set of 103 differential response genes to glucose (red dot).**

For each resampling, 103 genes were chosen at random from the genome. eSNPs for each gene were generated from GTEx (Version 7) using all 48 tissues. P-values for each eSNP were determined in our prior meta-GWAS for diabetic retinopathy*. The x-axis shows the proportion of eSNPs in each set with a FDR < 0.05 in the diabetic retinopathy meta-GWAS. The figure reveals a significant shift to the right (represented by the red dot) for the glucose response gene eSNPs in the meta-GWAS compared to resampled eSNPs.

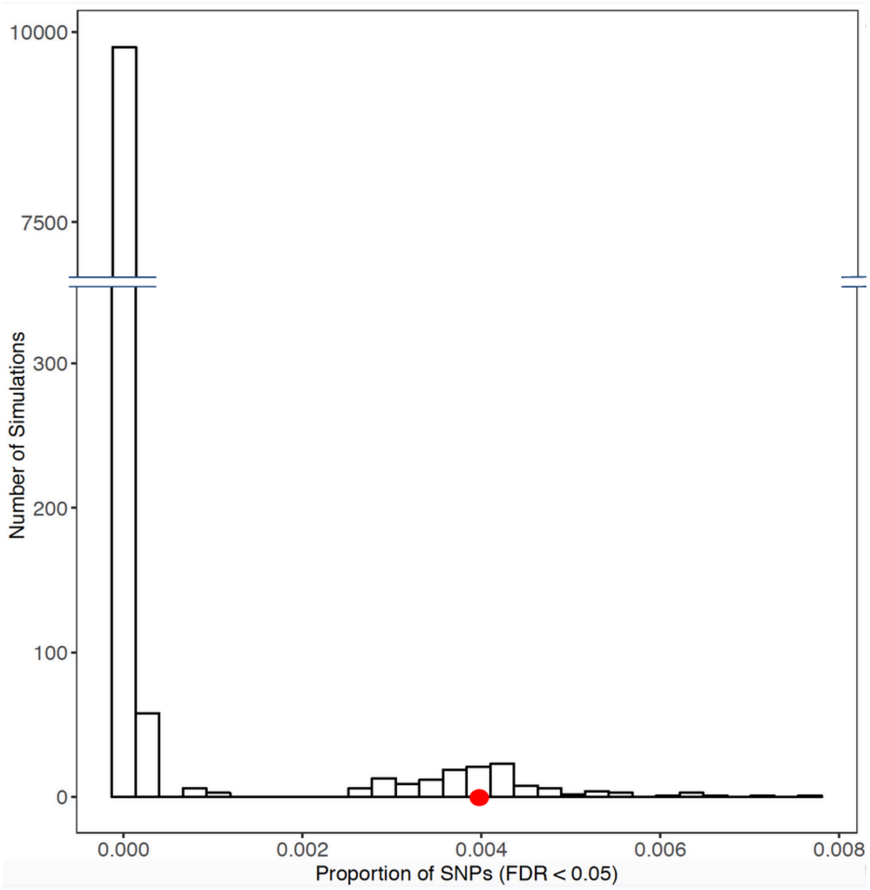



***** Grassi, M.A., et al., *Genome-wide meta-analysis for severe diabetic retinopathy.* Human molecular genetics, 2011. **20**(12): p. 2472-81.

**Fig. S7. FLCN expression in the human retina.**

(**A**) FLCN (green) is evident in the ganglion cell layer (GCL), neuronal cells of the inner nuclear layer (INL), and faintly in the outer nuclear layer (ONL). (**B**) Colocalization of FLCN with CD31 (red), a marker of endothelial cells, confirms FLCN expression in retinal blood vessels (white arrows).

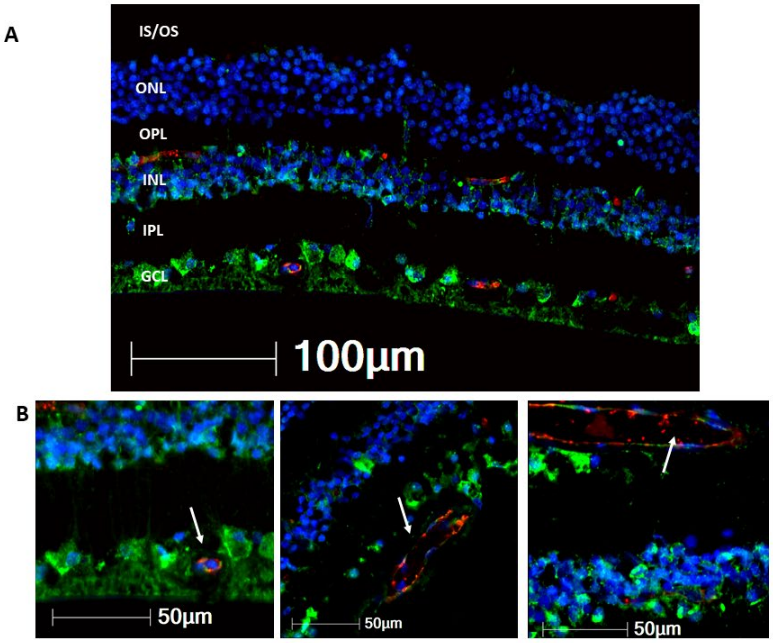

**Fig. S8. *FLCN* expression response to glucose by disease status (PDR vs nDR).**

Box and whisker plot of the change in *FLCN* expression in lymphoblastoid cell lines between standard glucose and high glucose conditions. Pink indicates the distribution of responses for individuals with proliferative diabetic retinopathy (PDR) ($log_2FC=0.08$), and blue indicates the same for individuals with diabetes but no retinopathy (nDR) ($log_2FC=-0.19$). Y-axis measures the difference in expression fold change on the $log_2$ scale. Each individual is represented by a dot. Difference in means between PDR and nDR is 0.27, p-value 0.003.

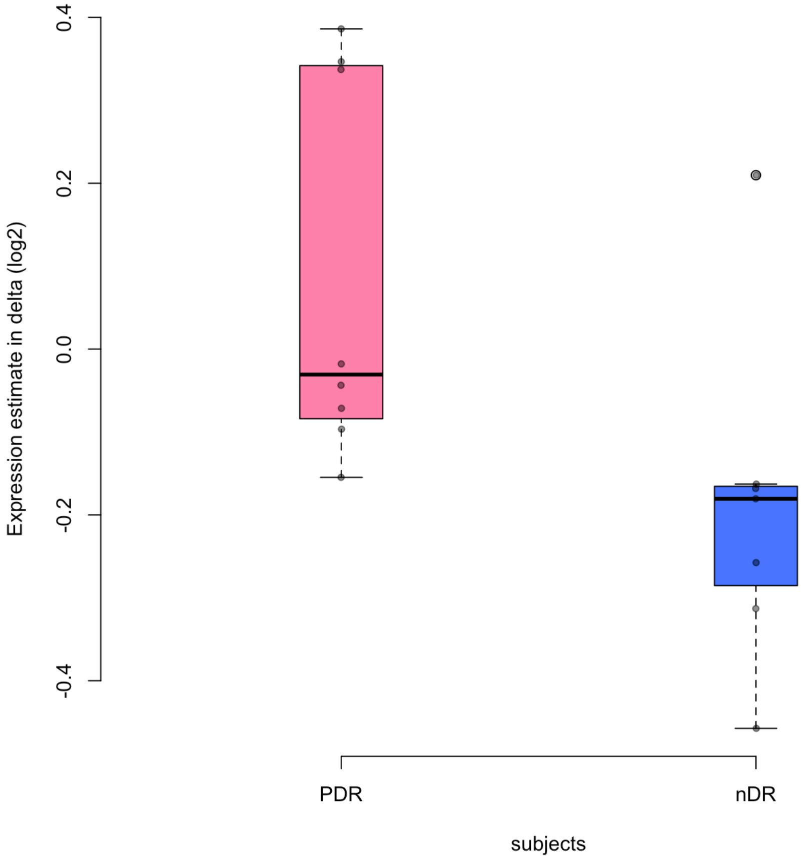

**Fig. S9. QQ Plot of diabetic retinopathy meta-GWAS p-values corresponding to 272 FLCN eSNPs.**

Points represent observed and expected GWAS-meta-analysis p-values for each of 272 *FLCN* eSNPs identified in the retina and more than 20 GTEx tissues. The red line represents the null hypothesis of no difference between the observed and expected p-values.

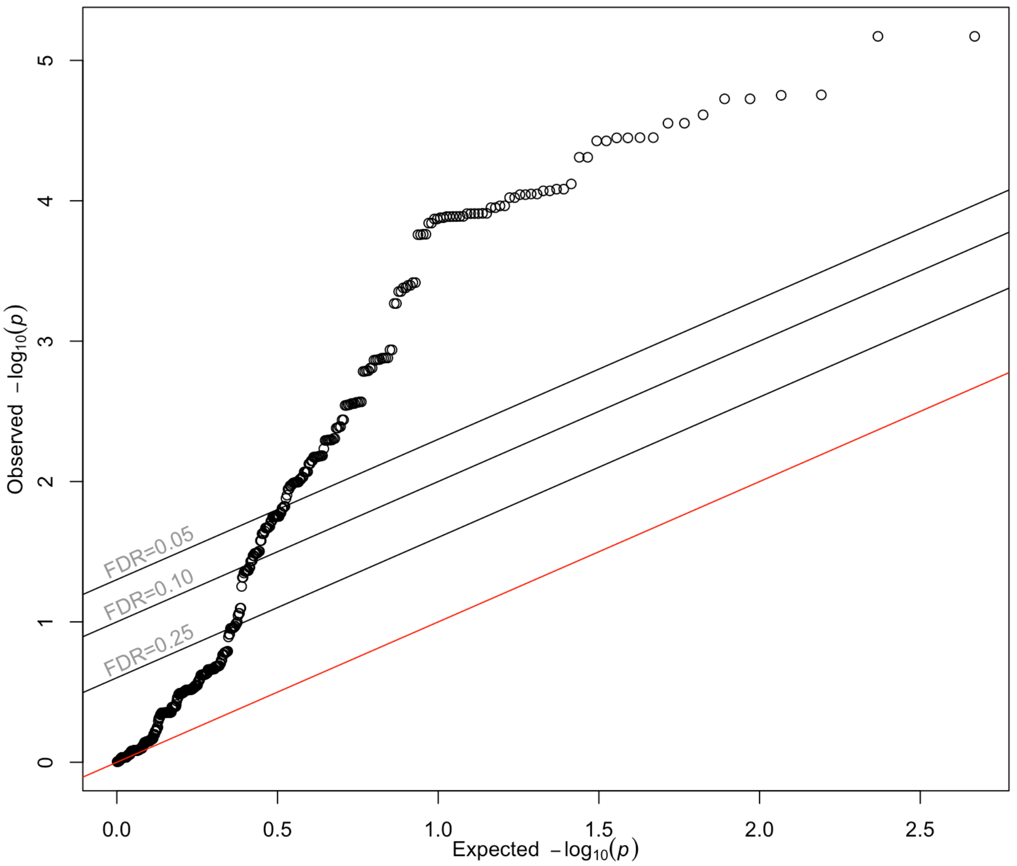

**Fig. S10. QQ Plot of UKBB diabetic retinopathy GWAS p-values corresponding to 272 FLCN eSNPs.**

Points represent observed and expected UKBB GWAS p-values for each of 272 *FLCN* eSNPs identified in the retina and more than 20 GTEx tissues. The red line represents the null hypothesis of no difference between the observed and expected p-values.

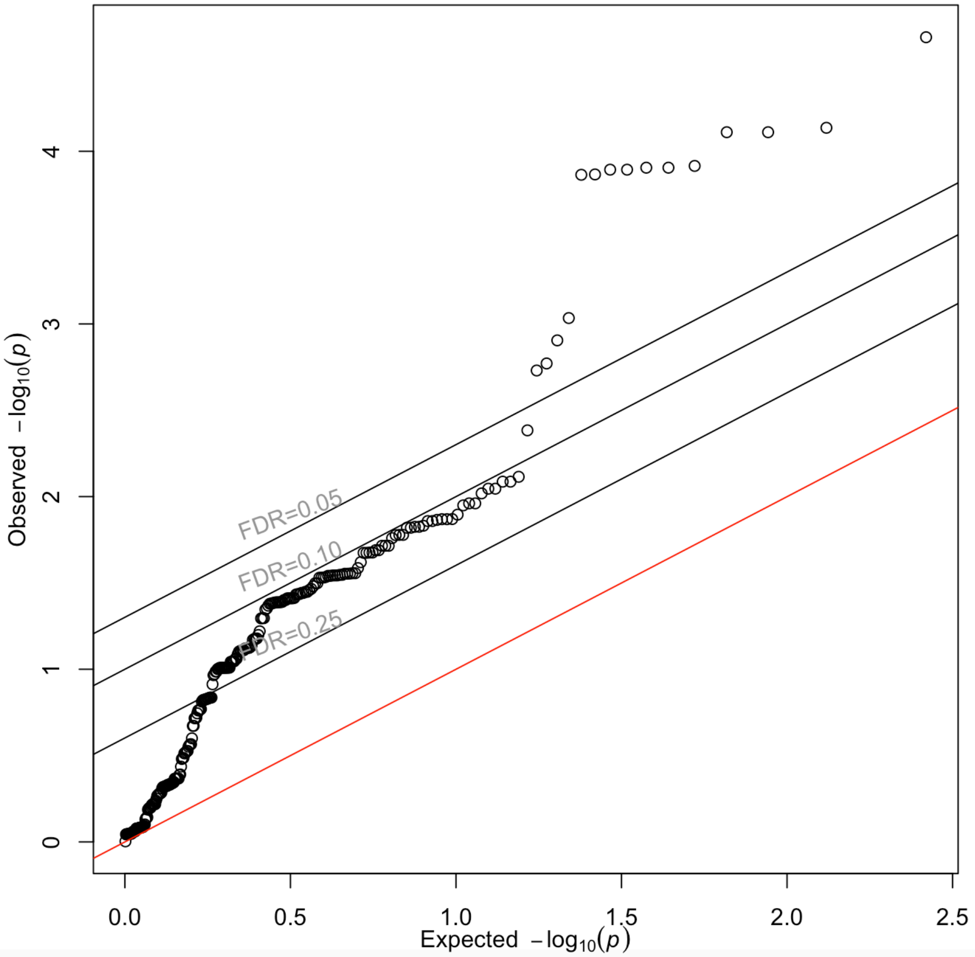
